# Modular adaptation of the ribcage in response to artificial selection for endurance running

**DOI:** 10.1101/2025.08.22.671715

**Authors:** Elizabeth Webb, Jesse Hennekam, Nicole E. Schwartz, Theodore Garland, Katrina Jones

## Abstract

The capability for sustained running has convergently evolved multiple times in mammals, and involves myriad anatomical, physiological, and behavioural adaptations. The ribcage plays a critical role in both respiration and locomotion but its adaptations to running are largely unexplored. Robustly testing adaptation in wild populations is challenging, so we use artificial selection for voluntary wheel-running behaviour (i.e., High Runner or HR mice) to directly test form-function relationships associated with sustained running. We compared ribcage configuration and shape of HR (males: 52, females: 47) to control (males: 48, females: 48) mice using rib counts and 3D Geometric Morphometrics. Two of four HR lines had an additional rib and increased variation in the proportion of true to false ribs, suggesting that ribcage patterning has been impacted by selection. This variability among lines suggests that selection for wheel running has resulted in adaptations that are expressed variably among the selected lines, resulting in “multiple solutions” to selection. Total ribcage shape did not vary significantly between HR and Control mice. Instead, the effect of selection varied along the ribcage, with significant effects in the caudal ribs. Further, the caudal ribcage of HR mice showed increased disparity, within-rib, and among-rib integration compared to controls. The strong response of caudal ribs indicates a modular pattern of adaptation, with cranial ribs possibly constrained by their role in ventilation. This study demonstrates that adaptation in the mammalian ribcage is likely shaped by a complex interplay of selection and craniocaudal integration and may result in variation at multiple anatomical levels (count, shape, modularity).

## Introduction

Cursoriality, specialization for fast or sustained running, has evolved multiple times across the mammal clade (Janis and Wilhelm, 1993; Lovegrove and Mowoe, 2014; Yuanqing et al., 2007), indicating a recurring pattern of selection for increasing running speed and/or endurance. Running adaptations have been linked with competitive predator-prey dynamics (Dawkins et al., 1997; Vermeij, 1993) and also with long-distance movement of ungulates due to changing environmental conditions in the late Miocene (Janis, 1993; Janis et al., 2004; Janis and Wilhelm, 1993). However, identifying cursorial adaptations in wild animals (e.g. Garland Jr and Janis, 1993; Kelly et al., 2006) can be challenging because disentangling locomotor adaptations from those related to other types of selection or from traits simply inherited from ancestors (“phylogenetic effects”) is complex (Garland Jr et al., 2005, 1993; Janis and Wilhelm, 1993; Rezende and Diniz-Filho, 2012). As an alternative to comparisons of wild species, artificial selection can be used to address such confounding factors by controlling for other aspects of variation among populations under known types of selection (Garland Jr and Rose, 2009). Although adaptations of the limb skeleton to cursoriality are well explored among species of mammals, little is known about how the axial skeleton may have adapted. In particular, the mammalian ribcage plays a critical role in ventilation and locomotion, and has been implicated in cursorial adaptations (Bramble, 1989). Here, we used mice artificially selected for high voluntary wheel-running behaviour to examine the potential relationships between running behaviour and ribcage morphology.

The High Runner (HR) mouse experiment started in 1993 (Swallow et al., 1998), with 224 mice from an outbred population of Hsd:ICR mice. After two generations of random mating, mice were split randomly into eight lines, four non-selected Control lines and four selected HR lines. Since then, once mice reach sexual maturity, they are given access to a wheel for six days. In the HR lines, the male and female (from within each family) that complete the highest number of revolutions on days five and six are selected to breed with an opposite-sex individual from another family of the same line for each generation. A similar procedure is applied in the Control lines, except that males and females are chosen from each family without regard to wheel revolutions. By generation 10 of selection, HR mice were running 75% more revolutions per day than Controls (Swallow et al., 1998). Eventually, a selection limit averaging almost 3-fold more revolutions per day was reached between generation 17-27, depending upon line and sex (Careau et al., 2013). The increased running distance of the HR mice is attributable mainly to faster average (and maximum) speeds, rather than the amount of time spent running, especially for females (Garland Jr et al., 2011; Girard et al., 2001).

Many aspects of the biology of the HR mice have been studied previously. Apparent adaptations associated with selective breeding include increased endurance and VO_2_max during forced treadmill exercise (Meek et al., 2009; Rezende et al., 2006; Schwartz et al., 2023; Singleton and Garland Jr, 2019), changes to specific brain regions associated with locomotor behaviour (Kolb et al., 2013; Schmill et al., 2023), larger hearts (Copes et al., 2015; Schwartz and Garland Jr, 2024), and larger femoral heads (Castro and Garland Jr., 2018; Garland Jr and Freeman, 2005; Kelly et al., 2006). In addition, the four replicate HR lines also differ from one another in morphology, physiology, and wheel-running behaviour (Garland Jr et al., 2011; Hillis and Garland Jr, 2023). Sex-specific differences have been observed in the response of the HR lines to selective breeding, including for daily running distance and speed (Careau et al., 2013; Garland Jr et al., 2011), organ masses (Swallow et al., 2005), and body fat composition (Hiramatsu and Garland Jr, 2018). Despite this extensive literature, including multiple studies on the appendicular skeleton of the HR mice (Castro and Garland Jr., 2018) and wild cursors (García-Esponda and Candela, 2010; Garland Jr and Janis, 1993), nothing is known about possible modifications of the thoracic cage (or ribcage) in the HR lines of mice.

Increased sustained running speed in mammals is associated with increased oxygen demands of muscles (Xu and Rhodes, 1999), and relatively increased maximal oxygen consumption (VO_2_max) has been observed in some cursorial mammals (Dlugosz et al., 2013; Lindstedt et al., 1991; Poole and Erickson, 2011). Movements of the thoracic cage are vital in generating lung ventilation (Carpenter, 1946; Hildebrand, 1974), although the relationship between ribcage morphology and ventilatory function is poorly understood. Furthermore, regardless of any training effects, HR mice have increased VO_2_max compared to Controls across multiple generations (Cadney et al., 2021; Schwartz et al., 2023; Singleton and Garland Jr, 2019; Swallow et al., 1998). In mammals more generally, increased VO_2_max has been correlated to increases in thoracic cage volume through rib shape change (Havryk et al., 2002; Weinstein, 2008) and changes in the ventilatory mechanism of cursors has been linked to thoracic shape change (Callison et al., 2019). Therefore, morphological changes in the ribcage of the HR mice could contribute to their increased VO_2_max.

In mammals, the thoracic cage is made up of the rib-bearing vertebrae and ribs, which together form the thoracic cage that supports the lungs (Flower and Gadow, 1885; Marlin et al., 2002). The true ribs articulate directly with the sternum whilst the false and floating ribs have no direct articulation to the sternum (Flower and Gadow, 1885). These more caudal thoracic vertebra form part of the diaphragmatic region, which is involved in running. Movements of the caudal-most, rib-bearing vertebrae have been recorded during asymmetrical running gaits of small mammal (Schilling and Hackert, 2006). Therefore, responses in the caudal ribcage might indicate adaptations that enhance axial running movements. We hypothesise that the ribcage of selectively bred HR lines may have modifications that could increase thoracic volume or spinal mobility during voluntary wheel running relative to the non-selected Control lines.

Here we examine the ribcage morphology of HR and Control mice, considering both ribcage composition and rib shape. Our hypotheses are as follows:

1. **Ribcage composition:** We expect increased rib count in HR mice to elongate the ribcage, and decreased within-line variability in rib count due to stabilizing selection that is associated with fast-running species(Galis et al., 2014; Gunji et al., 2025).
2. **Rib shape:** We expect individual rib shape and overall ribcage shape to differ between HR and Control lines. In HR mice, ribs may be longer and less curved to reflect the barrel shape observed in many cursorial species (Bramble, 1989; Hildebrand, 1974).
3. **Craniocaudal variation:** The influence of selection may vary craniocaudally along the ribcage. Further, we predict that craniocaudal variation in disparity, within-rib, and among-rib integration patterns will reflect modularity of function and evolvability of the ribcage (Lande, 1980; Olson and Miller, 1951).

## Methods

### Sample

The High Runner (HR) mouse experiment is an ongoing artificial selection experiment in which mice are bred for voluntary wheel-running behaviour (Careau et al., 2013; Swallow et al., 1998). The experiment was started using the genetically diverse Hsd:ICR strain (Carter et al., 1999). 224 mice from this strain were randomly bred for 2 generations and were then separated into eight closed lines, each consisting of at least 10 breeding pairs per generation. Pups were weaned at 21 days and then housed until ∼6-8 weeks of age in same-sex groups of four. At this point, individuals were separated and given access to a wheel for six days. For the four selected HR lines, selection was based on the number of wheel revolutions recorded on the 5th and 6th days. Control lines were subject to the same conditions, but breeders were chosen without regard to wheel running (Swallow et al., 1998). To reduce inbreeding, sibling mating was not allowed. Females and males were co-housed for 19 days with *ad lib* water and food (Harlan Teklad Laboratory Rodent Diet [W]-8604). Pregnant dams were then provided with a breeder diet (Harland Teklad Laboratory Breeder Diet [S-2335]-7004) until offspring weaning (at 21 days of age).

The mice used here were retired breeders from generation 93. Males were on average euthanized at 95 days old (after males were removed from breeding pairs, see above) and females at 126 days (one week after pups were weaned). Body mass was taken immediately after euthanization (via carbon dioxide inhalation). Then, body composition was measured using non-invasive quantitative magnetic resonance (EchoMRI-100; Echo Medical Systems LLC, Houston, Texas, USA), which independently calculates fat and lean mass. Body length was measured to the nearest mm with a ruler. Body composition and body length were repeated twice, and never in direct succession, to reduce measurement error, and subsequently averaged. Analyses of body mass and length were performed using SAS Procedure MIXED, with line nested within linetype as a random effect (e.g. see Castro and Garland Jr., 2018; Copes et al., 2015; Hiramatsu and Garland, 2018; Schwartz et al., 2023).

Carcasses were frozen at −20 C and then µCT scanned postmortem at the University of Southern California using a Nikon XT225ST scanner at 80kv 120uA, 1440 projections producing scans with 61μm pixel size. Osteological material was segmented out of scans using Dragonfly ORS software version 2022.1.0.1259 (Comet Technologies Canada Inc, 2020). Osteological meshes were then converted to 3D surface wraps in meshlab version 2022.02 (Cignoni et al., 2008), using Poisson surface reconstruction (Kazhdan and Hoppe, 2013), to reduce time taken for further analysis. In total 100 male and 95 female mice were used (Table 1).

**Table 1:** Sample size for Control and High Runner mice. F: Female; M: Male.

| Linetype | Control |  |  |  |  |  |  |  | High Runner |  |  |  |  |  |  |  |
| --- | --- | --- | --- | --- | --- | --- | --- | --- | --- | --- | --- | --- | --- | --- | --- | --- |
| Line | 1 |  | 2 |  | 4 |  | 5 |  | 3 |  | 6 |  | 7 |  | 8 |  |
| Sex | F | M | F | M | F | M | F | M | F | M | F | M | F | M | F | M |
| Number of Specimens | 12 | 12 | 11 | 12 | 12 | 12 | 13 | 12 | 12 | 12 | 11 | 16 | 12 | 12 | 12 | 12 |

### Rib count and ribcage composition

The typical ribcage composition was: 13 ribs total with 7 true and 6 combined false and floating ribs; this matched the typical mouse thoracic and vertebral formula reported in the literature (Asher et al., 2011). True ribs were defined as ribs attached directly to the sternum via a single cartilage, false ribs articulate to the sternum via a shared cartilage, and floating ribs do not articulate to the sternum (Hildebrand, 1974). When a rib was present on only one side of the individual it was counted as half thoracic and half lumbar, e.g., producing 13.5 thoracic vertebrae. For analysis, false and floating ribs were grouped together to reduce the number of possible combinations of rib types. Meristic variation was defined as variation in the overall total number of ribs. Homeotic variation was defined as changes, relative to the typical composition, that involve addition of one rib type and loss of an adjacent rib type (Cerbus et al., 2024). Ribcage compositions were plotted by linetype (Control or HR) and replicate line (1, 2, 4 and 5 being Controls and 3, 6, 7, and 8 being HR). SAS Procedure GLIMMIX was used to identify significant differences in both the frequency of typical and atypical ribcage composition and the typical total rib count (13) and atypical total rib count. These were tested by replicate line and linetype to assess statistical significance. Initial analyses also included sex and the sex-by-line or sex-by-linetype interaction terms, but these were found to be nonsignificant and so were dropped from the final models.

### Rib shape

Rib morphology was quantified using 3D geometric morphometrics. Landmarks were placed using 3D Slicer software, version 5.0.2 (Kikinis et al., 2014). A total of 33 homologous landmarks were placed per rib (Figure 1 and Table S1), including 8 anatomical and 25 semi-landmarks along two curves characterising the interior surface (12 landmarks) and exterior surface (13 landmarks). The landmarking protocol was adapted to capture the length and curvature of the rib (Bastir et al., 2017; García-Martínez et al., 2018). The articular tubercle was not present on all ribs. To capture this feature, a redundant landmark was placed (Jones et al., 2018; Klingenberg, 2008a) at the nearest equivalent point, which was the caudal point of the rib head.

**Figure 1:**
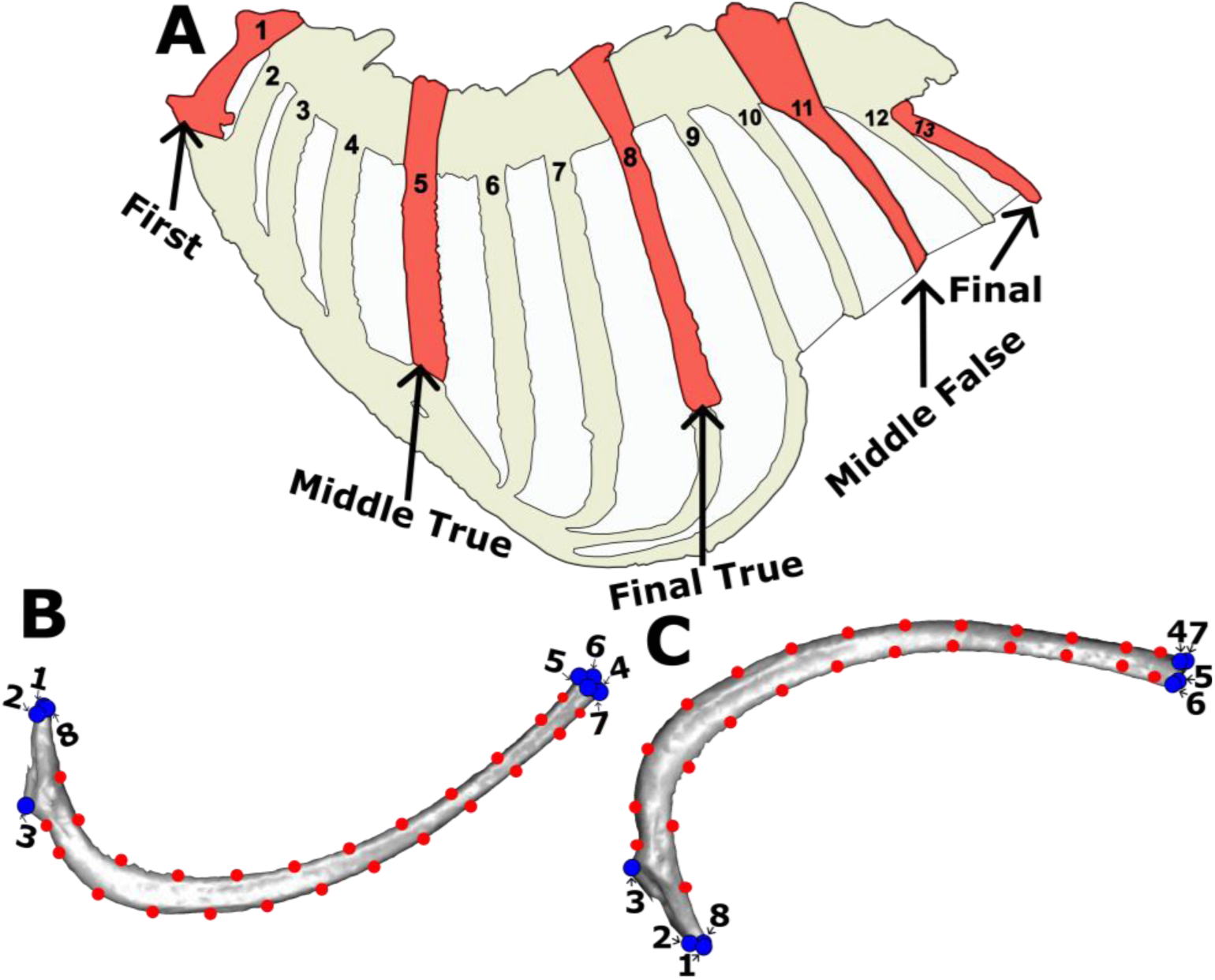
Landmarking scheme. A. Positions of ribs selected for analysis. Trace of specimen 96181. Rib 7 from specimen 96063 shown in caudal (B) and cranial (C) view. Blue: fixed landmarks, Red: sliding semi-landmarks. Landmark descriptions can be found in Supplemental Table S1.

To ensure the landmarking approach was repeatable, a sensitivity analysis was conducted on pilot data collected from each rib along the left side of the ribcage of six specimens (Supplementary Information). From these data (Figure S1), we chose five presumed-homologous rib positions along the rib cage that captured the major craniocaudal variation represented in the sample. These were the first rib, the middle true rib, the last true rib, the middle false/floating, and the last rib. If the rib count was an even number, the more caudal middle rib was always used; for example, if there were 8 true ribs, middle rib 5 was always chosen (rather than middle rib 4). The sensitivity analysis was repeated with these five ribs to ensure that it still captured a range of morphological variation (Supplementary Information Methods and Figure S1 and S2). The sexes were analysed separately as females were older than males at time of sacrifice, had undergone pregnancy and parturition, and exhibit significant sexual dimorphism in skeletal traits associated with the hindlimb (Castro and Garland Jr., 2018). Effect of line was tested as a nested variable. In addition, the effect of line was also tested per linetype (i.e., effect of line was tested separately for Controls and HR) and gave generally similar results (Tables S5-7).

Shape analysis was conducted using the R package geomorph (version 4.0.5, Baken et al., 2021). A generalised Procrustes superimposition was done (GPA), using the function *‘gpagen’*. This aligns the landmarks in the same shape space and removes the effects of translation, rotation, and size (Rohlf and Slice, 1990; Zelditch and Swiderski, 2023). The 23 semi-landmarks were allowed to slide along their respective curves during the Procrustes fit. A separate Procrustes fit was performed for each of the five rib positions separately. To visualize the morphological variation in individual ribs, a principal component analysis (PCA) was conducted using *‘gm.prcomp’* in geomorph for each of the five rib positions. We used a male Control Line 1 specimen (Specimen: 96206) without any obvious vertebral anomalies for rib shape warping. Due to the relative subtlety of the shape variation, maxima and minima associated with PC1 and 2 were magnified by 3.

To examine total ribcage shape, we ran a new Procrustes fit for all the ribs from all the specimens together and plotted them in a single combined PCA. To account for serial variation, these Procrustes coordinates were then concatenated for the five ribs into one set of coordinates per individual mouse (Jones et al., 2018). This analysis treats each landmark on each rib as a different variable from the same individual, instead of the same variable on multiple individuals (ribs). This allows visualization of the variation that is common across the serial repeating structures, in this case the ribs. To visualize shape variation across multiple ribs simultaneously using 3D meshes, concatenated coordinates for each rib were translated in space to visualize them individually. Each rib was then rescaled using the mean centroid size associated with that rib position (code available via https://figshare.com/s/0eda5951131c004f6f09). The rib mesh object was then warped to the minimum and maximum concatenated shape of both PC1 and PC2. A few apparent outliers were detected visually and original models were rechecked to ensure there were no landmarking errors (see Results).

The impact of selective breeding on individual and concatenated rib shape was tested using a nested ANCOVA. Procrustes coordinates were the dependent variable and compared with Line nested within Linetype. Body length was included as a covariate to account for allometry (Klingenberg, 2016), as body length is a better predictor of skeletal morphology than body mass in the HR mice (Castro et al., 2020). All models were run using *‘procD.lm’* and were tested using Type III sums of squares and 10,000 iterations. Line was nested within Linetype as each line represents an independent repeat of the experiment. To account for this, the error effects tested in *‘procD.lm’* were adjusted using the function *‘anova’.* The effect of Linetype was judged relative to the error associated with line, instead of the error associated with the residuals.

### Craniocaudal variation

To examine craniocaudal variation in relation to selective breeding, the relative effect of line and linetype at each rib position was assessed using the Z score generated from the individual rib ANCOVAs. The Z score is calculated as the standard deviation of the F value of the given model relative to all the permuted null models, and is a good estimator of effect size in complex models (Collyer et al., 2015). We also tested craniocaudal variation in morphological disparity (Guillerme et al., 2020; Hopkins and Gerber, 2017) and within-rib integration (Klingenberg, 2008b) at each rib position. Morphological disparity was calculated from the Procrustes variance for each rib position using the function *‘morphol.disp’*. Within-rib integration was calculated by measuring covariation in eigenvalues at each rib position (Conaway and Adams, 2022) using the function ‘*integration.Vrel’*. Among-rib integration patterns were also measured to assess modularity in the ribcage. Between-rib integration was measured pairwise between each rib position using a singular warps analysis in the function *‘integration.test’*. As a heuristic, differences between HR and Control lines in ribcage integration were then compared using a two-tailed t-test on the r-PLS scores in both males and females, and between males and females, using the function *‘t.test’*.

## Results

### Body Length and Mass

For males, HR and Control lines did not significantly differ for body length (SAS Procedure Mixed with REML estimation and Type III tests of fixed effects: N = 100, F_linetype_ = 1.12, d.f. = 1,6, P = 0.3314; Least Squares Means and associated Standard Errors were 10.79 ± 0.08 mm for Control and 10.67 ± 0.08 for HR lines). Results were similar for females (N = 95, F_linetype_ = 1.46, P = 0.2720; LSM ± SE were 10.98 ± 0.08 mm for Control and 10.83 ± 0.08 for HR lines). For males, HR mice tended to weigh less than Control mice (N = 100, F = 5.49, P = 0.0576; LSM ± SE were 34.08 ± 1.34 g for Control and 29.66 ± 1.33 for HR lines), but this trend was less apparent for females (N = 95, F = 1.51, P = 0.2649; LSM ± SE were 34.36 ± 1.18 g for Control and 32.31 ± 1.18 for HR lines). We also analysed body mass with length as a covariate. For males, length-adjusted mass was statistically lower for HR mice (F = 6.42, P = 0.0444; LSM ± SE were 33.59 ± 1.00 g for Control and 30.01 ± 1.00 for HR lines), but not for females (F = 1.30, P = 0.2976; LSM ± SE were 34.03 ± 0.85 g for Control and 32.65 ± 0.85 for HR lines). Considering variation among the replicate lines, body length significantly varied for HR males (N = 52, FF_line_ = 12.04, d.f. = 3,48, P < 0.0001), HR females (N = 47, F = 6.03, d.f. = 3,43, P = 0.0016), Control males (N = 48, F = 2.83, d.f. = 3,44, P = 0.0491), and Control females (N = 48, F = 4.74, d.f. = 3,44, P = 0.0060).

### Hypothesis 1: Rib count and ribcage composition

The typical rib count was 13, consisting of 7 true ribs and 6 combined false and floating ribs. Variation was observed in the total number of ribs (13, 13.5 or 14) and in ribcage composition (8T 5F, 7.5T 6.5F, 8T 5.5F, 8T 6F, 7.5T 5.5F, 5T 8F, 7T 6.5F, 6T 7F, 6T 8F, 7T 7F). Non-typical rib cage composition was observed in all but one line (HR Line 3), with non-typical conditions involving the gain of a mixture of false and true ribs (Figure 2). In most cases, increases in the total number of ribs were associated with gain of false ribs (although 8T 6F configurations were observed in HR lines 8 and 6, see Figure 2). Most lines exhibit gain of both true and false ribs. However, HR Lines 8 and 6 have a large proportion of individuals with additional false ribs due to increased thoracic count in HRs (Table 2, Rib count: p=0.0130). The overall percentage of non-typical composition in Control lines varied from 2% to 5%, while in the HR lines it varied from none in Line 3 to 87.5% in Line 8 (Table 2). The frequency of non-typical ribcage composition varied significantly among lines (p=0.0008); specifically, HR lines 6 and 8 had many more non-typical compositions than in any Control line (Figure 2). GLIMMIX analysis revealed there was no significant difference between the HR and Control lines once line is taken into account for count or composition (p=0.4314/0.4108), but line was highly significant (p=0.013/<0.001).

**Figure 2:**
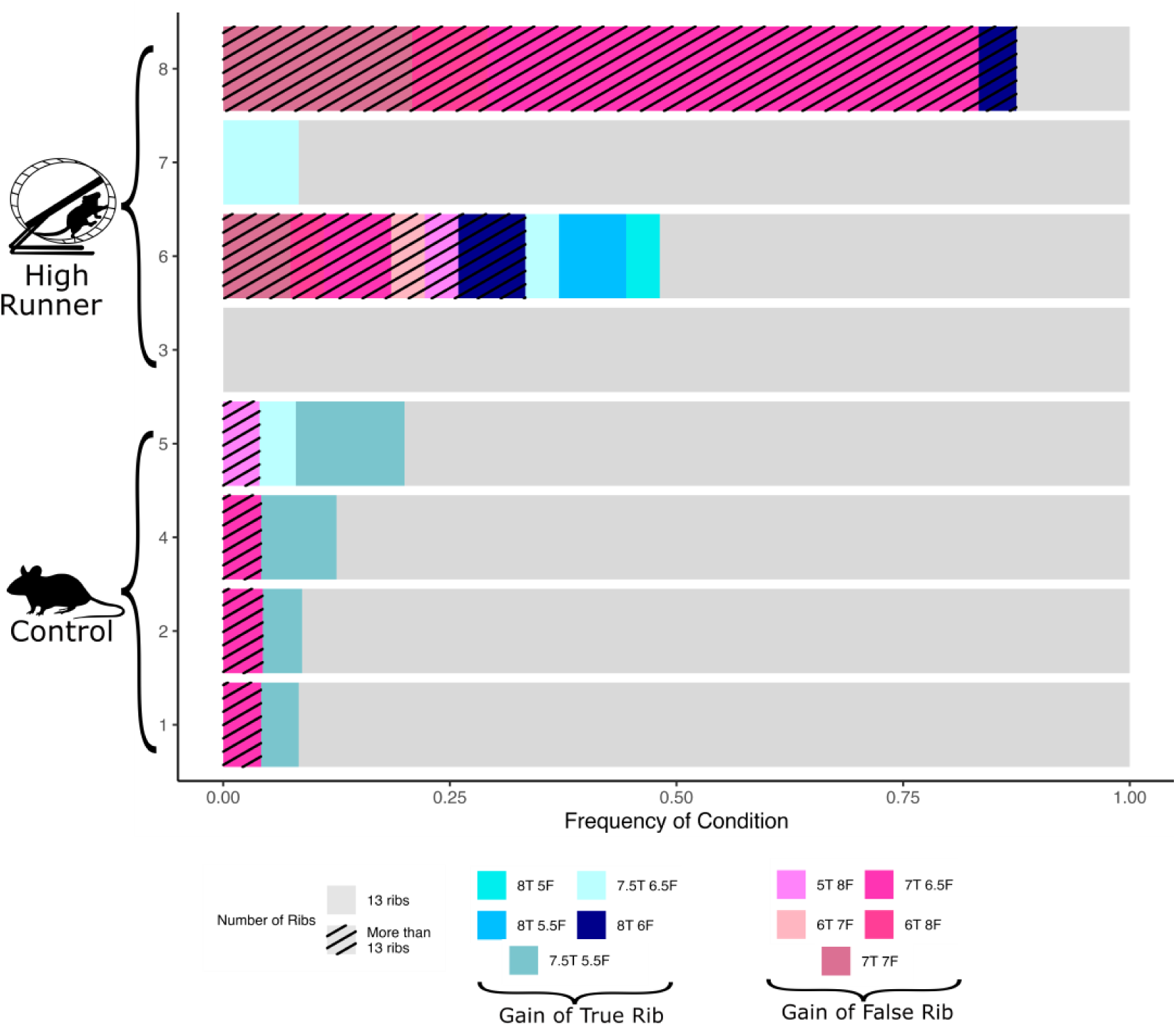
Ribcage composition in selectively bred High Runner and replicate Control lines. Pattern indicates rib counts and colour indicates rib compositions. Sample includes both males and females (total N = 195).

**Table 2:** SAS Procedure GLIMMIX analysis of ribcage composition and thoracic count.

|  |  | Ribcage Composition |  | Rib Count |  |
| --- | --- | --- | --- | --- | --- |
|  |  | Typical | Non-typical | 13 | ≠13 |
| **Control** | **1** | 92% (22) | 8% (2) | 96%(23) | 4% (1) |
|  | **2** | 91% (21) | 9% (2) | 96%(22) | 4% (1) |
|  | **4** | 88% (21) | 13% (3) | 96% (23) | 4% (1) |
|  | **5** | 80% (20) | 20% (5) | 96%(24) | 4% (1) |
| **HR** | **3** | 100% (24) | 0% (0) | 100% (24) | 0% (0) |
|  | **6** | 52%(14) | 48%(13) | 67% (18) | 33% (9) |
|  | **7** | 92% (22) | 8% (2) | 100% (24) | 0% (0) |
|  | **8** | 12.5%(3) | 88% (21) | 13% (3) | 88% (21) |
| **P (Line)** |  | <0.001 |  | 0.0130 |  |
| **P (Linetype)** |  | 0.4108 |  | 0.4314 |  |

### Hypothesis 2: Rib shape

Figure 3 shows that individual lines tend to cluster in morphospace, and while some lines are quite distinct from others, there is considerable overlap between mice from the HR and Control lines. In addition, Control lines typically have more overlapping morphospace than HR lines. Most variation in rib morphology is observed in HR Line 7 of the true ribs of females and HR Line 8 at the caudal end of the ribcage in males and females to a lesser extent (Figure 3). For Line 7 of true ribs of females, there was increased rib curvature and increased length of the rib head. At the middle false and final rib of Line 8, ribs were straighter and thinner in females and straighter but thicker in males compared to other HR and Control lines

**Figure 3:**
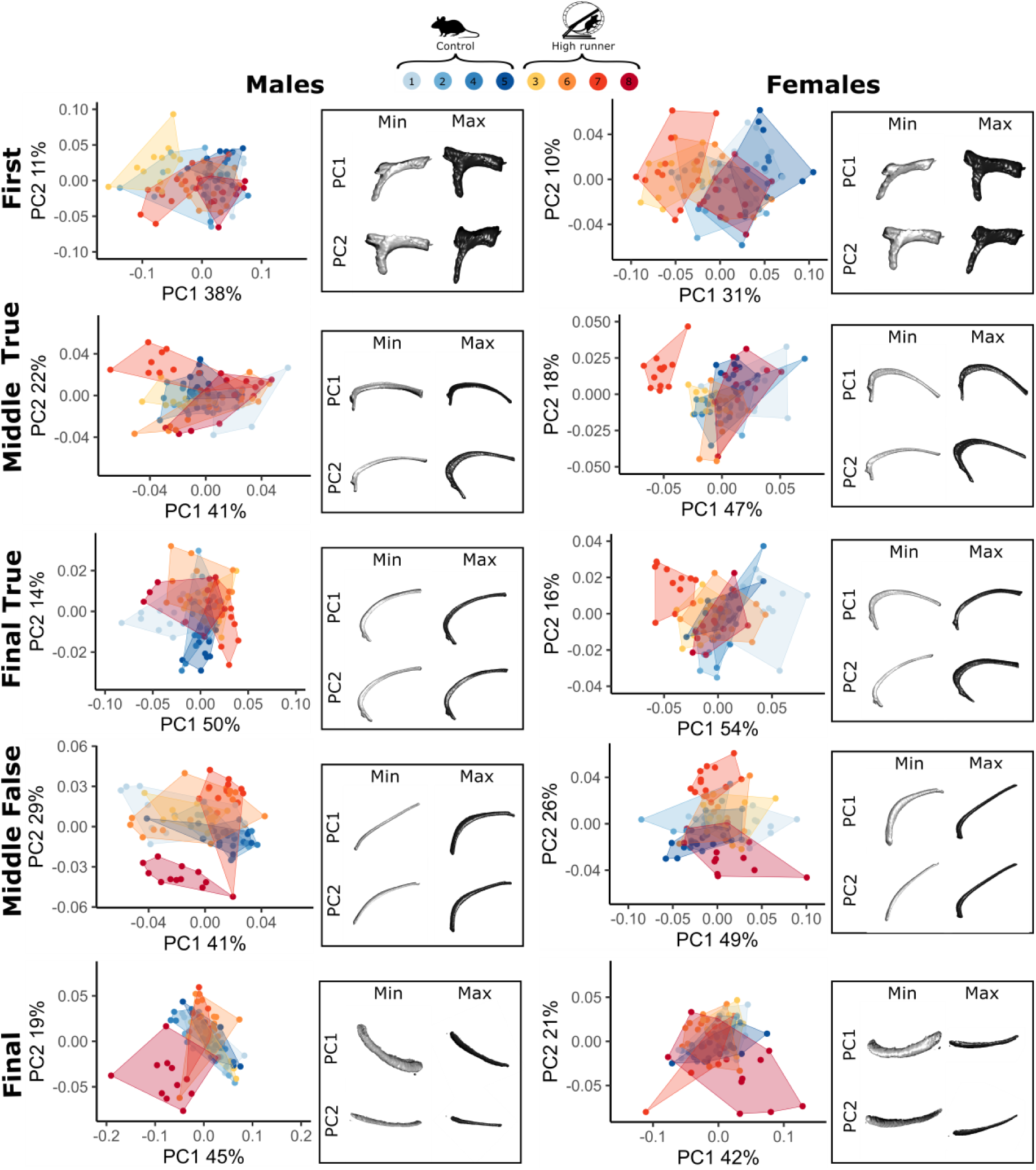
PCA of shape variation at each rib position separated by sex for PC1 and 2. Black boxes show extremes of shape variation along each PC, magnified by three (see Methods). Specimens that appeared to be outliers because they deviate from their line specific cluster were checked for scan quality and landmarking accuracy before inclusion (e.g. outlier in F3 bottom right panel). Axes are scaled to maximise the variation.

Rib variation at each position of the ribcage is visualised along PC1 and PC2 (Figure 3). Together PC1 and 2 accounted for between 41% and 75% of total variation at each rib position (Figure 3). At the first rib, PC1 represents variation in rib curvature, while PC2 reflects variation in rib length. For the middle and final true rib, PC1 captured increasing curvature, while PC2 captured thickening of the rib. The middle false rib exhibited the most variation in rib curvature, with negative PC1 representing a relatively straight rib, and positive PC1 indicating curvature in the middle for males and nearer the head for females. The middle false rib of females also had a narrowing of the rib at the maximum of PC1. The major variation in the finals ribs was in the relative thickness of the rib, with both males and females becoming thinner along PC1 and flatter along PC2.

When all the ribs are plotted in a single morphospace, the major axis of variation is the craniocaudal position of the ribs, with PC1 accounting for 75% of the variance (Fig. 4A). This confirms that craniocaudal patterning is the dominating signal when comparing overall variation in rib shape. The total ribcage shape was analysed by concatenating the individual ribs into a single landmark configuration that accounts for craniocaudal variation in each specimen. In males, PC1-4 each accounted for >5% of variation and together contributed to 47% of variation, with PC1 and PC2 contributing 19% and 11% (Fig. 4C). In females, PC1 to 5 each individually accounted for >5% of variation and together contributed to ∼47% of variation, with PC1 and 2 accounting for 19% and 14% of variation respectively (Fig. 4B). PC3 and PC4 for the concatenated dataset can be found in Supplemental Figure S3 only PC3 and 4 were visualised as further PCs represent less than 5% of variation. Shape variation associated with the concatenated PC axes can be found in Supplemental Video 1.

**Figure 4:**
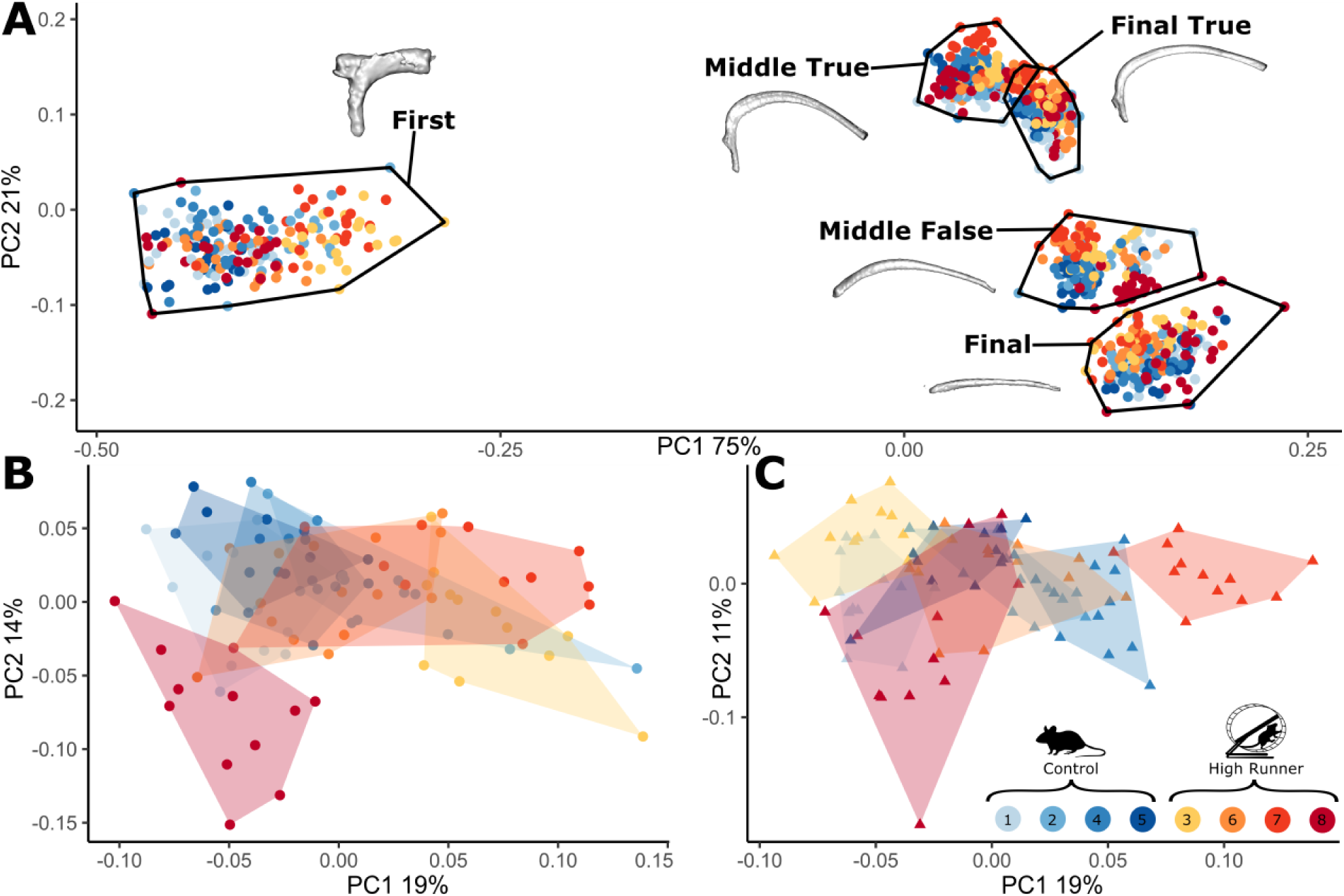
**A** PCA along the entire ribcage. Most variation is due to differences by rib position. Rib shapes are mean shapes associated with each rib position. **B** Males and **C** Females, PC1 and PC2 of rib shape concatenated across five rib positions.

As with the individual rib analysis, morphospace overlap between HR and Control mice is considerable, but varies by line. In both males and females, HR Line 6 tends to cluster strongly with the Controls, while HR Lines 8, 7, and 3 have more distinctive morphology (which additionally differs by sex). Line 8 tends to have more negative PC1 and PC2 scores in males and lower PC2 scores in females. This may reflect the high frequency of additional ribs in this line. Line 7 has more positive PC1 scores, particularly in females, which relates to high curvature in the true ribs. Line 3 is characterized by high PC1 scores in males, but low PC1 scores in females, and seems to represent a relatively straight first rib. Thus, each of these three lines seem to have modified a different portion of the ribcage in response to selection (Line 3: first rib, Line 7: true ribs, Line 8: false ribs). There were limited differences by line along PC3 and PC4 of males with much of the morphospace overlapping (Supplemental Figure S3).

For individual rib shape, Procrustes ANCOVA revealed that the greatest impact was from replicate line, which was significant (p=<0.05) at each rib position for both males and females (Table S4 and S5, Fig 5). Similar patterns were also seen when separated by linetype (Control shape∼line+length and HR shape∼line+length) with line being significant in all positions in females and in the first three ribs in males (Table S6-S9). By contrast, the impact of linetype on rib shape was only significant at one position for females and males (final true rib, p=0.037; final false, p=0.007 respectively) and it was marginally significant for males at the middle false position (p=0.059). For the concatenated rib shape, ANCOVA on concatenated Procrustes coordinates showed highly significant effects of replicate lines (nested within linetype) in both males and females, but no overall effect of Linetype (Table 3). This suggests that although there are some significant differences in rib shape at specific points along the ribcage between the selectively bred HR lines and their non-selected Control counterparts, the overall morphological variation in ribcage shape is primarily driven by differences among the replicate lines within the two treatment groups.

**Figure 5:**
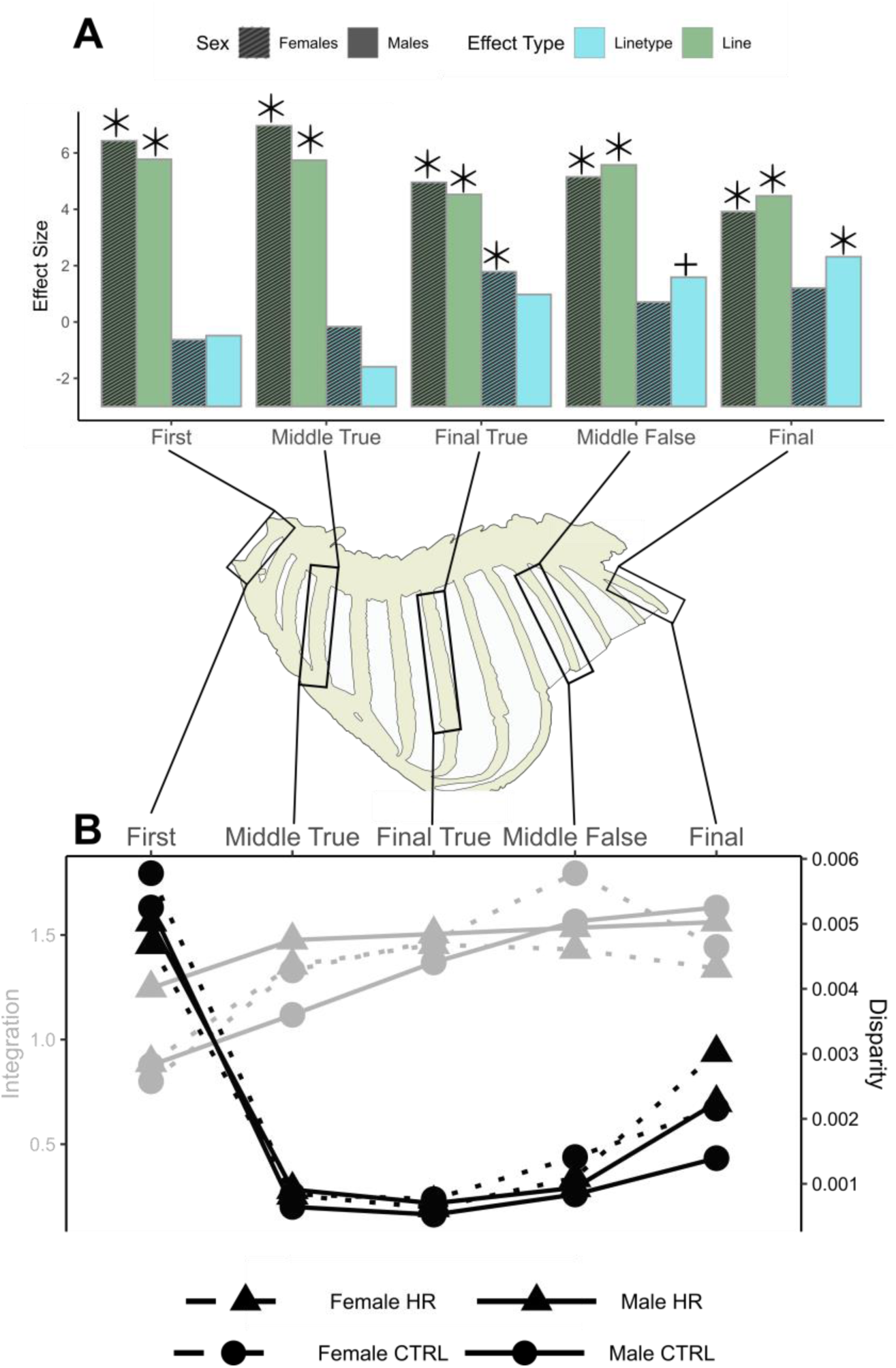
Shape variation by rib position in relation to linetype and line separated by sex. **A**. Effect size calculated as Z value from MANCOVA analysis (using formula shape∼linetype/line for each rib position for separate sexes) in ProcD.lm (Table S4 and S5). Star indicates statistical significance at 0.05 level and + indicates statistical significance at the 0.1 level. **B**. Within-rib integration estimated using relative eigenvalue index and disparity calculated using the Procrustes variance.

**Table 3:** Results of ANCOVA of concatenated Procrustes coordinates separated by sex using Type III Sums of Squares to match the design of the HR mouse selection experiment (see Methods).

|  | Male |  |  |  |  |  |  |
| --- | --- | --- | --- | --- | --- | --- | --- |
|  | Df | SS | MS | Rsqr | F | Z | Pr(>F) |
| **Linetype** | 1 | 0.06801 | 0.06801 | 0.04656 | 1.11611 | 1.24944 | 0.10999 |
| **Body length** | 1 | 0.00959 | 0.00959 | 0.00656 | 0.88262 | -0.22939 | 0.58414 |
| **Line(nested)** | 6 | 0.36562 | 0.06094 | 0.25030 | 5.60926 | 9.29677 | 0.00010 |
| **Residuals** | 91 | 0.98858 | 0.01086 | 0.67677 |  |  |  |
| **Total** | 99 | 1.46073 |  |  |  |  |  |
|  | Female |  |  |  |  |  |  |
|  | Df | SS | MS | Rsqr | F | Z | Pr(>F) |
| **Linetype** | 1 | 0.05009 | 0.05009 | 0.03793 | 0.92622 | 0.74470 | 0.22788 |
| **Body length** | 1 | 0.00968 | 0.00968 | 0.00733 | 0.95086 | -0.02272 | 0.51155 |
| **Line(nested)** | 6 | 0.32450 | 0.05408 | 0.24572 | 5.31129 | 10.63760 | 0.00010 |
| **Residuals** | 86 | 0.87572 | 0.01018 | 0.66312 |  |  |  |
| **Total** | 94 | 1.32060 |  |  |  |  |  |

### Hypothesis 3: Craniocaudal variation

Comparing effect sizes and significances from the ANCOVAs along the ribcage revealed variation in the response to selection. The effect of line was significant in males and females for all rib positions. The effect size of linetype (HR vs Control) was considerably smaller than that of line but tended to be higher in the last three rib positions for both males and females (Figure 5 and Table S4 and S5). In males, the effect of linetype was particularly strong in the last two rib positions.

Morphological disparity differed significantly between all rib positions (all p values<0.001). Morphological disparity was greatest at the first rib but then decreased drastically at the middle true rib before gradually increasing caudally until the final rib for Control and HR males and Females (Figure 5). This pattern was consistent across males and females for both HR and Control individuals. Morphological disparity was higher in HR males and females at the first and final rib position compared to Controls (Table S10 and S11). There were significant differences between within rib integration values in the first rib however other rib positions were not significant (Table S12 and S13). Significant differences by sex and linetype in Within-rib integration were observed in the first rib and less commonly in the middle true rib For Control females, integration increased up to the middle false rib, where it then decreased. For HR females, integration increased up from the first to the final true rib, where it decreased. However, for Control and HR males, integration increased along the length of the ribcage. Selected males had higher integration in the true ribs than in Control males.

Integration between different rib positions varied by sex and linetype. Compared to Controls, integration values for HR individuals were significantly higher for females (t= 2.1584, p= 0.0454) but not males (t= 1.1506, p= 0.2649). There was also no significant difference between female and male integration values (t= −0.77104, p= 0.4455). In Control mice, integration tended to be less with the final and middle false rib, suggesting that the true ribs may form a distinct module (Figure 6). However, selection appears to have impacted this pattern. While integration values are still higher in the true ribs, the more caudal end of the ribcage is more integrated in HR mice compared to Controls. Specifically, there is no significant integration between the middle false and final rib in Control males or females. However, the middle false and final rib are significantly integrated in both males and females from the HR lines.

**Figure 6:**
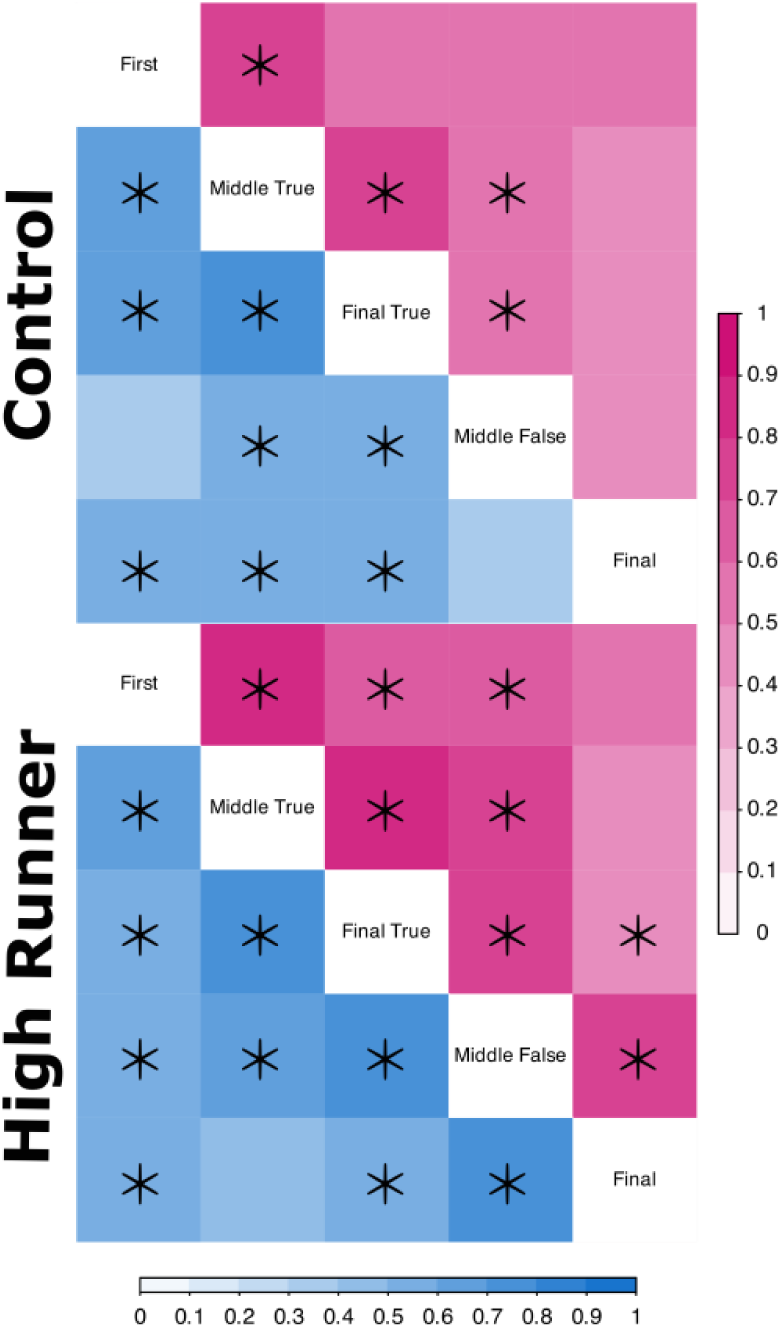
Integration between different rib positions by sex and linetype. Darker colours indicate higher levels of integration between ribs. Pink indicates integration for females and blue indicates integration for males. Significant integration at the p=0.05 or below are marked with *.

## Discussion

The thoracic cage is a critical component of the mammalian ventilatory apparatus that functions to support the lungs and assist the diaphragm during breathing. Despite this, studies examining morphological variation in the mammalian ribcage are rare relative to other anatomical regions, such as skulls and limbs (Callison et al., 2019). The HR lines of mice have been selectively bred for increased voluntary wheel--running behaviour, which we hypothesized may have resulted in changes to the ribcage morphology that increase thoracic volume and facilitate their elevated maximal rates of oxygen consumption. Here, we have explored morphological variation in the four replicate HR and four replicate Control lines of mice and demonstrate diverse modifications of the thoracic region in conjunction with long-term selective breeding for voluntary wheel running. Of the four selected lines, two (HR 6 and 8) have evolved changes in rib counts and ribcage composition, while three lines (HR line 3, 7 and 8) have evolved modified rib shape. Impacts of selective breeding were strongest in the caudal ribs, and together with disparity and integration patterns, suggest a ‘diaphragmatic’ module in the ribcage that is more influenced by locomotor adaption and less constrained by ventilatory function.

### Some HR lines have homeotic increases in rib count relative to Control lines

The morphology of the mammalian ribcage is determined by the number of ribs, the relative proportion of different rib types (ribcage composition), and the shape of the ribs. Although some data exist on total rib counts (Li et al., 2023) we have little information on variation in ribcage composition in mammals. This is because different rib types are defined by the nature of their cartilaginous attachments to the sternum that are rarely preserved in skeletal preparations in museum collections. However, using CT scans of intact cadaveric specimens, we were able to identify true false and floating ribs based on their intact position. The high degree of variation in relative counts of true and false ribs highlights ribcage composition as an important and evolutionarily labile aspect of ribcage morphology.

Two of the four HR lines showed both an increase in overall rib number and a change in ribcage composition (HR line 6 and 8). Both lines typically had a higher number of individuals, with atypical rib counts (33% of HR line 6 and 87% of HR line 8 vs 12% of other Control and HR lines) indicating a longer ribcage relative to the other lines. They also vary in their ribcage composition. HR line 6 had increased numbers of all rib types, while HR line 8 only had increased numbers of false and floating ribs. The role of different rib types specifically in mammalian ventilation is unclear (Cieri et al., 2018), making functional interpretations challenging. However, increasing the length of the ribcage could be an adaptation for increasing thoracic volume and facilitating higher maximal oxygen consumption associated with increased running ability. Although HR lines 3 and 7 generally did not have alterations to their rib number or composition, they may possess other adaptations related to their higher maximal oxygen consumption and running ability (i.e., multiple solutions, as described in Garland Jr et al., 2011; Hillis and Garland Jr, 2023). Clearly, this is an area worthy of further study, both within and among other species of mammals.

In the vertebral column, homeotic variation is defined as the repatterning of one vertebral or rib type to another adjacent type, whereas meristic variation is defined as the loss or addition of a vertebra or rib (Bateson, 1894). In the present study, increases in rib number were always accompanied by a reduction in the number of adjacent lumbar vertebrae, thus indicating that these were homeotic changes in the thoracolumbar boundary. Previous work has suggested that cursorial mammals may have a constraint on variation in total thoracolumbar count compared with non-cursorial relatives (Galis et al., 2014; Williams et al., 2019), due to the functional demands of running. However, we find the opposite, that lines of mice bred for high endurance-running behaviour have elevated variation in craniocaudal patterning. Functional constraints on vertebral counts may be limited to the lumbo-sacral boundary, thus only impacting thoracolumbar count in cursorial mammals. By contrast, homeotic shifts between thoracic and lumbar identities seem to be less subject to functional constraints and even show increased variability under selection for running. Homeotic changes in rib patterning are controlled by temporal and/or spatial shifts in hox code patterning in the presomitic mesoderm (Cerbus et al., 2024; Christ et al., 2007; Mulley, 2022). The total number of ribs is determined by the expression of *Hox10* as this patterns the thoracolumbar boundary (Carapuço et al., 2005; Mallo et al., 2010), while the number of ribs that connect to the sternum (true and false ribs but not floating) is controlled by *Hox9* (McIntyre et al., 2007). However, as the sternum is derived from appendicular tissues, the lateral plate mesoderm (LPM) (Brent et al., 2023), the true-false/floating rib boundary may also be influenced by changes in expression of the LPM. Evolutionary-developmental mechanisms are rarely considered in studies of behavioural evolution (Hoke et al., 2019). It has been shown that *HoxB* has been altered in selected mice (Hillis et al., 2024) and *HoxB* is involved in early development of the presacral region including the thoracic region (Wilmerding et al., 2021). Therefore, further study of the hox mechanism in the HR mice may be a fruitful avenue for future research.

Previous work on earlier generations of HR mice hypothesized that increased inbreeding, and subsequent deterioration of traits (inbreeding depression), may have impacted skeletal morphology in both Control and selected HR lines (Castro et al., 2021). Increased inbreeding in combination with artificial selection in domestic dogs, has produced changes in vertebral formula (Brocal et al., 2018). Further, cheetahs have undergone extensive inbreeding in their evolutionary history long after their adaptations for fast running (Dobrynin et al., 2015; Menotti-Raymond and O’Brien, 1993). Despite this, cheetahs have low variability in thoracolumbar count, indicating that strong natural selection related to functional constraints may limit inbreeding depression in cursorial species (Galis et al., 2014). Mammals have a strongly conserved vertebral formula, with rodents having one of the most conserved vertebral formulas of all mammalian groups (Asher et al., 2011; Li et al., 2023; Narita and Kuratani, 2005). Both Control and HR lines show increased rib count variability compared to wildtype lab mouse populations (Asher et al., 2011). However, changes in rib count in Control lines of mice are at relatively low frequency compared to the high levels observed in the two selected HR lines (Line 6 and 8), suggesting an effect beyond inbreeding depression and possibly adaptive.

### The impact of selection on rib shape is focused at the caudal-most ribs

Ribcage shape can impact thoracic volume and thus could influence ventilatory function and tidal volume. For example, cursorial mammals tend to have barrel-shaped ribcage (Bramble, 1989; Hildebrand, 1974) and have increased tidal volume during exercise (Callison et al., 2019). Similarly, humans living at high altitudes have increased rib length and decreased rib curvature, traits that presumably facilitate VO_2_max in low oxygen availability environments (Little and Haas, 1989; Weinstein, 2008). HR mice have increased VO_2_max relative to Controls, however, this physiological adaptation for endurance running is not accompanied by expected morphological adaptations in rib shape. Overall, the impact of selection for voluntary wheel running on rib shape was subtle, with the effect size (measured as the Z score) of HR vs Control being 0.75-1.25 in overall ribcage shape and −1.6-2.3 of total variation in individual ribs, as compared with effect size of 9.2-10.6 for line of whole ribcage shape and 6.96-3.96 for line by individual ribs (Table 3, S4 and Figure 5). We found no statistically significant effect of linetype on overall ribcage shape, nor on the shape of the first two rib positions (first and middle true ribs). However, there was a significant or marginally significant effect on the last three positions (female: last true, male: middle and last false), where effect sizes were consistently higher, indicating some response to selection in the caudal ribcage.

We predicted that ribs of HR mice might be longer and less curved, thus producing a more barrel-shaped chest, as is observed in many cursorial mammal species (Bramble, 1989; Hildebrand, 1974). For example, it has been hypothesised that the deep and narrow ribcage adopted by fast-running species allows for the scalene and the sclenatomastoid muscles to be coopted into running (Bramble, 1989; Hildebrand, 1974). However, we did not find such changes in HR lines. Shape variation associated with selection is very subtle in these lines of mice and seems to relate to shortening of the ribs, though some straightening is observed in the middle false position. One possible explanation for this is that morphological variation may in some cases reflect the homeotic shifts in false ribs described above. For example, the addition of a false rib in HR Line 8 results in a final rib that tends to be smaller than in the individuals with a typical rib count. As the ribs extend down the cranio-caudal length of the ribcage they become flatter (Figure 3). Therefore, homeotic shifts may have similar impacts that propagate up the ribcage, demonstrating the complex relationship between serial variation among ribs and that among individuals.

By contrast, we found a strong effect of line in both overall ribcage shape and at every individual rib position, for both sexes. When experiencing the same type of selection, populations will often adapt somewhat differently, and this process can be facilitated by random genetic drift (e.g., see discussion in Lande, 1976). Differences among the replicate HR lines have been demonstrated with respect to various traits (Castro et al., 2024; Schwartz et al., 2023; Whitehead et al., 2023), including how they have evolved increased daily wheel-running distance, with variation in how much they have increased the duration of running versus average running speeds (Garland Jr et al., 2011). In the present study, some lines appeared more divergent in their morphology, and these tended to be HR lines (Figure 3 and Figure 4B and 4C). Morphological differences were present in both shape and position along the ribcage. For example, in HR line 7 females they have more divergent true ribs, which typically reflect increased curvature, whilst HR line 8 has straighter false ribs in both sexes, and HR line 3 has less curved first ribs. This indicates that natural diversity may produce different modifications to the ribcage under the same selection pressures.

Several factors may influence rib shape in the HR lines of mice. Glucocorticoids (e.g., cortisol, corticosterone) have a wide range of effects on bone (Martin et al., 2021). HR mice have higher circulating corticosterone levels than Controls (Girard and Garland Jr, 2002; Malisch et al., 2007; Singleton and Garland Jr, 2019). Therefore, it is possible that shape changes seen in the ribs of certain lines are a non-adaptive by-product of high corticosterone levels seen across all HR lines, although more so for some lines than others (e.g., see Malisch et al. 2007). Or alternatively, the increased corticosterone levels may have facilitated adaptive changes in ribcage shape.

Additionally, the mini-muscle MM phenotype observed in some HR mice (HR lines 3 and 6 Garland et al. 2002) may be influencing rib shape. Some individuals from HR line three, in which the MM phenotype is fixed, are in a distinct morphospace from other lines (Figure 3 and 4) and there were no abnormal ribcage compositions observed in HR line 3 (Figure 2). HR line 3 is genetically different from all other HR and Control lines (Hillis and Garland Jr, 2023). All MM individuals show various increased organ masses compared to non-MM individuals (Rezende et al., 2006; Swallow et al., 2005). Rib shape changes seen in MM individuals might be a consequence of increased sizes of internal organs, as opposed to being an adaptive response that increases ventilatory capacity or running ability. However, the MM phenotype is also present, but is not fixed, in HR Line 6 (where such differences were not as apparent). Further investigation into MM individuals in Line 3 and Line 6 are needed to see if similar effects on the ribcage of this phenotype can be replicated.

The impact of both linetype and line varied between males and females, suggesting an interaction between selection, genetic drift, and sex. The function of the ribcage is dimorphic in humans (García-Martínez et al., 2019). Many sex differences have been noted in morphological traits of the HR mice (e.g. see Castro and Garland Jr., 2018) and these may be contributing to running behaviour differences between the sexes (Garland Jr et al., 2011). It is also possible that pregnancy, parturition, and their more advanced age has affected female ribcage morphology relative to males. Indeed, the ribcage changes morphology during pregnancy in humans (LoMauro et al., 2019), although the long-term effects of this after pregnancy are unknown. Furthermore, ribcage morphology changes throughout aging and the males in this study are much younger than the females, which this could also be impacting ribcage morphology (Gayzik et al., 2008; Weaver et al., 2014). Therefore, the sex differences seen in the rib shape could be related to differing responses to selection, age, or pregnancy.

### Craniocaudal variation in disparity and integration patterns suggests modularity in adaptive responses

Morphological disparity is the degree of shape variation at a single rib position and was greatest in the first rib across the lines and sexes (Figure 3 and 5). The first rib is developmentally different from the others, as it attaches directly to the manubrium and not the sternebrae at the sternal end. Here, developing somitic (primaxial) and lateral plate (abaxial) components interact, and so this region is referred to as the lateral somitic frontier (Durland et al., 2008; Mitchel et al., 2022; Nowicki et al., 2003). The lateral somitic frontier is thought to pose a developmental constraint on the first rib (Buchholtz, 2014; Shearman and Burke, 2009) owing to a need for precise interaction between these two disparate tissues. Despite this, we see very high morphological disparity at this position. This result suggests that the composite nature of the first rib may in fact promote evolutionary lability, enabling it to respond to the dual influences of the axial and appendicular skeletal development. This idea should be tested at an interspecific level.

Within-rib integration patterns also varied along the ribcage. The rib is composed of several regions: the head, which articulates with the vertebral body, the neck, which articulates with the vertebral arch, and the rib shaft, which supports the intercostal muscles. Therefore, within-rib integration can shed light on the degree to which these different structures are varying in a modular versus coordinated way. In vertebrae, high within-element integration correlates with high disparity and ecology, suggesting that integration patterns promote evolvability (Jones et al., 2018). However, integration could also limit disparity when the morphological variation is misaligned with the selective regime (Felice et al., 2018). In the present study, within-rib integration correlates well with apparent variation in the selective regime, as the ribs with highest integration are also those with the strongest effect of Linetype. Similarly, if we exclude the first rib, there is also some degree of correlation with disparity. Disparity increases caudally from the middle true to the final rib, particularly in males. Thus, if we consider there may be a distinct developmental mechanism driving the extremely high disparity at the first rib, then we have evidence for a craniocaudal trend in the response to selection, within-rib integration, and disparity. This suggests that the caudal ribcage may form a more-evolvable region, associated with running (Schilling and Hackert, 2006), distinct from the more cranial ‘true ribs’.

We examined modularity of the ribcage by comparing the patterns of among-rib integration between HR and Control mice. Our results suggest that there is a modular pattern of variation in Control mice, with relatively high integration among true ribs, but limited integration between the false/floating ribs and the rest of the ribcage. Selection for voluntary wheel running has not only increased overall integration but has also promoted integration among the false ribs, e.g., between the final rib and middle false rib. It has been hypothesised that integration of the skull and jaw elements facilitates extreme feeding behaviours in snakes (Vincent et al., 2006) and that integration of limb bones facilitates the evolution of specific locomotor types (Villmoare et al., 2011) – but none have explored the potential role of integration in the ribcage (to our knowledge). Our data provides direct evidence that selection may increase integration within the ribcage. By contrast, data from the appendicular skeleton does not generally support the idea that selection promotes integration among limb bones in HR mice (Young et al., 2009), likely because the impacts of selection can vary across regions of the skeleton.

These patterns of integration reflect functional and developmental modularity within the ribcage. In humans, the upper (1^st^-5th ribs) and lower parts of the ribcage (6^th^-10^th^ ribs) develop and function separately from each other (Bastir et al., 2013; Okuno et al., 2019). It has been hypothesised that the upper portion of the ribcage, that is the true ribs, are more involved in mammalian ventilation (Bastir et al., 2017; De Troyer et al., 2005; Yoganandan and Pintar, 1998). But this is largely derived from *ex vivo* passive inflations of the lungs and has not been validated *in vivo.* However, in some reptiles, the caudal end of the ribcage is more involved in ventilation (Cieri et al., 2018) and in healthy humans it is seen that the caudal ribs rotate more during breathing (Luu et al., 2021). Further, the most caudal thoracic vertebrae are more actively involved in running than the more cranial ones, by contributing to sagittal bending of the spine during running gaits (Schilling and Hackert, 2006). These vertebrae are also more specialised in cursorial mammals, apparently reflecting locomotor adaptation (Jones et al., 2018). Therefore, the selective response in caudal ribs may reflect recruitment of the caudal ribs into a ‘locomotor module’ that involves locomotor functions associated with the lumbar spine.

In the vertebral column, the caudal-most thoracic vertebrae share a mix of features from the thoracic and lumbar regions and are thus considered transitional. At the ‘diaphragmatic’ vertebra, the vertebral zygapophyses transition from horizontal (thoracic-type) to vertical (lumbar-type) and the caudal-most vertebrae are therefore known as the post-diaphragmatic thoracics, or the diaphragmatic region. This region has been linked with locomotor adaptation, as well as high disparity and rates of evolution in mammals (Jones et al., 2018). The diaphragmatic vertebra in mice lies around T10, and thus falls within the false and floating ribs, suggesting a similar transitional region in the ribcage to that observed in the vertebral column. This may reflect the recruitment of the diaphragmatic region (ribs and vertebrae) into functions that characterize the lumbar region, and/or it may indicate that these caudal ribs have escaped from functional constraints associated with ventilation due to their lack of true articulations with the sternum.

