## Supplemental Information for "Modular adaptation of the ribcage in response to artificial selection for endurance running"

Table S1. *Anatomical positions of Landmarks.*

| ***Number*** | ***Anatomical Position*** | ***Landmark Type*** |
|---|---|---|
| ***1*** | *Cranial point of the rib head* | *Fixed* |
| ***2*** | *Caudal most point of the rib head* | *Fixed* |
| ***3*** | *Lateral point of the articular tubercule* | *Fixed* |
| ***4*** | *Cranial point of the sternal end* | *Fixed* |
| ***5*** | *Caudal most point of the sternal end* | *Fixed* |
| ***6*** | *Exterior medial point of the sternal end* | *Fixed* |
| ***7*** | *Interior medial point of the sternal end* | *Fixed* |
| ***8*** | *Medial distal point of rib shaft* | *Fixed* |
| ***9-21*** | *Exterior curve from points 9-21* | *Sliding* |
| ***22-33*** | *Interior curve from points 22-33* | *Sliding* |

*Sensitivity analysis: Methods*

To ensure the landmarking protocol could capture shape variation a sensitivity analysis was carried out. Six mice were chosen to display a range of linetypes, lines, sexes and thoracic counts to capture the largest range of ribcage morphologies (Table S2). The landmarking protocol was conducted on 13 thoracic ribs on the anatomical left-hand side of the mouse. The 14th rib was not included as it resembles the morphology of the 13th and by not including it we could concatenate individual rib shape data to form a whole ribcage dataset. Each mouse was landmarked three times in a randomised order with no reference to previous attempts. A total of 234 ribs were sampled and used for the sensitivity analysis.

*Table S2:* Metadata associated with the individuals chosen for the sensitivity analysis.

| Mouse ID | Linetype | Line | Sex | Thoracic Count |
|---|---|---|---|---|
| 96206 | C | 1 | M | 13.5 |
| 96037 | C | 2 | F | 13 |
| 96022 | C | 5 | F | 13 |
| 96530 | HR | 6 | M | 13.5 |
| 96471 | HR | 3 | M | 13 |
| 96365 | HR | 8 | F | 14 |

All shape analysis was conducted using the package geomorph (version 4.0.5, Baken et al., 2021) in R studio. Landmarks were subject to Procrustes superimposition to remove the effect of size, rotation and reflection using the function *'gpagen'*. This data was then subject to a principal component analysis (PCA). Procrustes coordinates from individual rib data were then concatenated to create a dataset to represent total ribcage shape. This dataset was then subject to a PCA (Jones 2018). A MANOVA was carried out on the concatenated procrustes coordinates using *'procDlm'* with individual mouse and rib number as factors.

We then plotted the individual rib shape variation and repeated the ANOVA with just the subsample data intended to be used for the final dataset (five ribs only), to ensure that the subsampling process was sufficient to still identify shape variation.

*Sensitivity analysis: Results*

We found that the largest shape difference was between the different ribs sampled along PC1 and PC2 (Figure S1A). The sub-sampling procedure also maintained this spread of shape variation across the morphospace (Figure S1B). Following concatenation shape data was plotted along PC1 and 2 (Figure S2A). Repeats were clustered in the shape space by individual mouse indicating that the landmarking protocol is accurate enough to be able to identify individual mouse morphology, the results of the MANOVA confirm this with no significant difference (p=0.526) in procrustes coordinates of repeats of the same mice (Table S3). The process was repeated with just the five ribs used for the final analysis. The repeats were still clustered and the morphospace is close to that of the original dataset (Figure S2B). These results were then confirmed by the MANOVA resulting in the same significance in the results (Table S3).

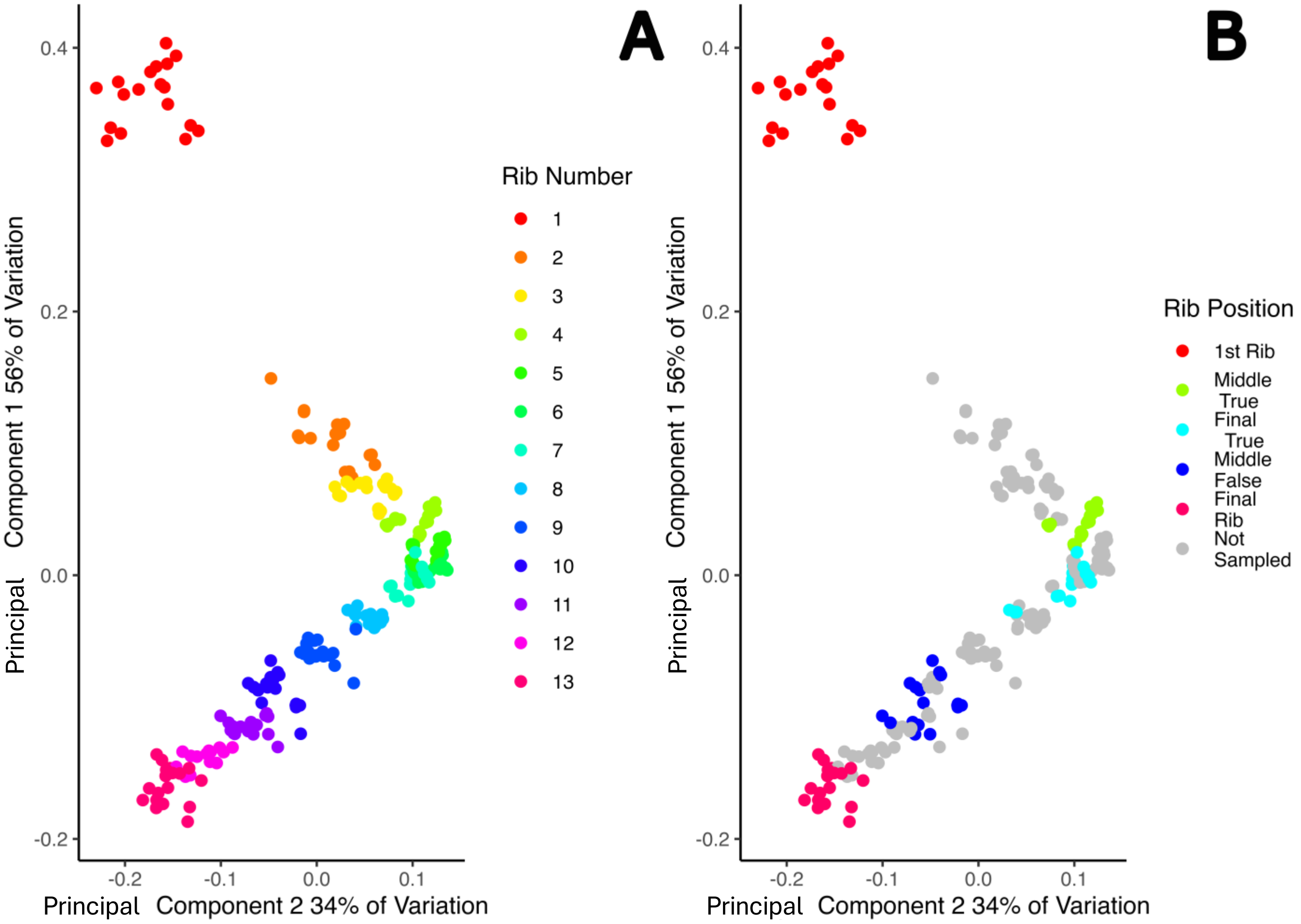


**Figure S1:** **A** Shape variation of individual ribs shown along principal component (PC) 1 and 2. Shape variation is determined predominantly by rib number. **B** Shape variation of the ribs that will be used for the final analysis sample. The selected sample still represents a wide range of morphospace

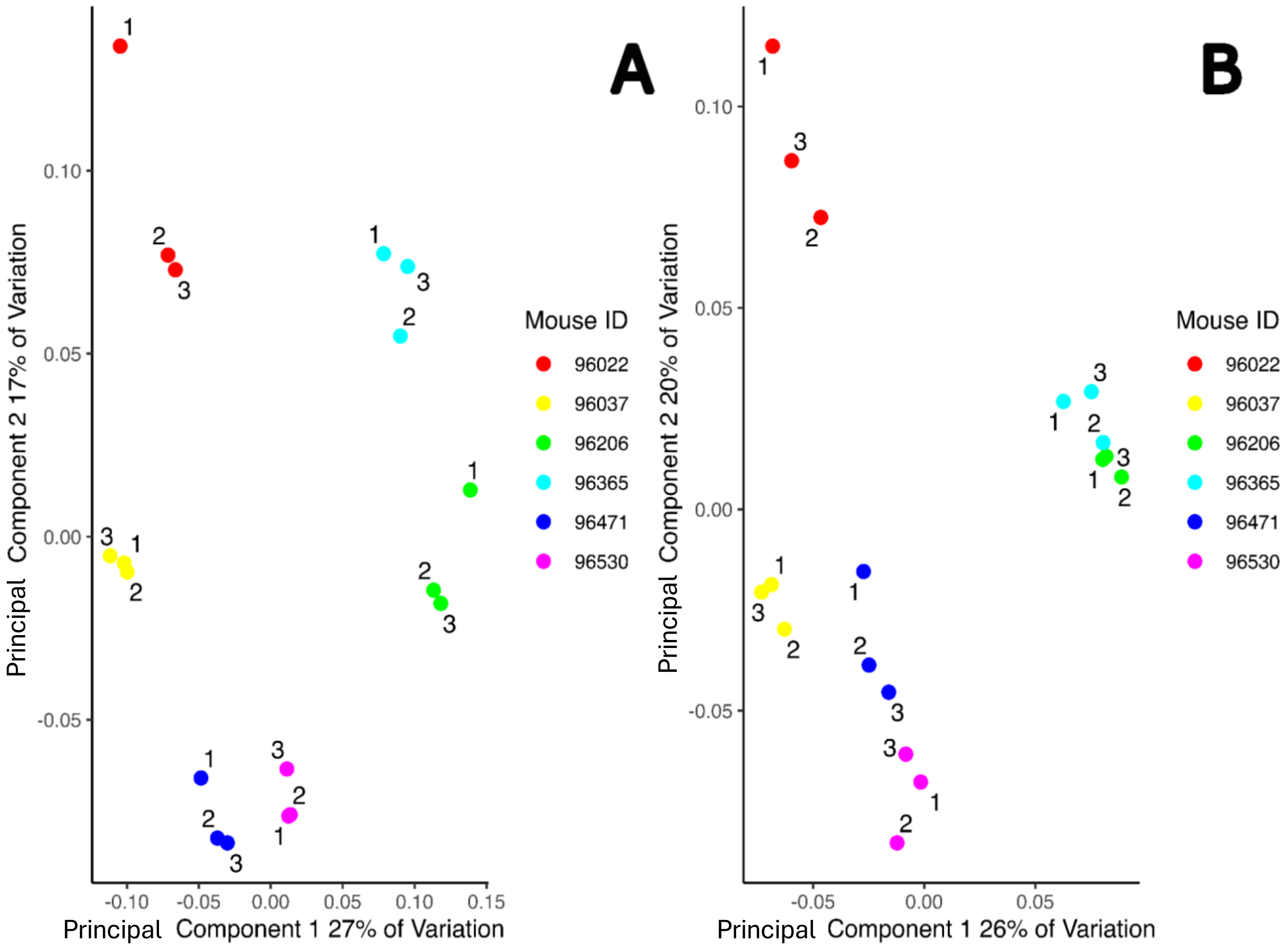


**Figure S2: A** Concatenated whole ribcage shape variation. 1,2,3 indicate the repeat number. Each repeat is clustered in the same shape space indicating that the landmarking protocol can detect shape variation. **B** Concatenated whole ribcage shape variation with only the five ribs to be used for the final analysis. Whilst the morphospace has changed with a marginal increase in principle component variation on PC1 and 2, repeats are still clustered by mouse ID showing that the subsampling process is good enough to be able to maintain shape variation

**Table S3**: Results of MANOVA analyses on all data and just the sample data to be used for final analyses

| **All Data** | | | | | | | |
|---|---|---|---|---|---|---|---|
| | **Df** | **SS** | **MS** | **Rsq** | **F** | **Z** | **P** |
| **Mouse** | 5 | 0.072957 | 0.014591 | 0.010192 | 0.922266 | -0.06741 | 0.526 |
| **Rib** | 1 | 3.493633 | 3.493633 | 0.488073 | 220.8192 | 6.944125 | 0.001 |

| | | | | | | | |
|---|---|---|---|---|---|---|---|
| **Residuals** | 227 | 3.591421 | 0.015821 | 0.501734 | | | |
| **Total** | 233 | 7.158011 | | | | | |
| **Sample Data Only** | | | | | | | |
| **Mouse** | 5 | 0.054731 | 0.010946 | 0.011556 | 0.300404 | -2.164555 | 0.984 |
| **Rib** | 1 | 1.657262 | 1.657262 | 0.349902 | 45.481616 | 4.782348 | 0.001 |
| **Residuals** | 83 | 3.024359 | 0.036438 | 0.638542 | | | |
| **Total** | 89 | 4.736351 | | | | | |

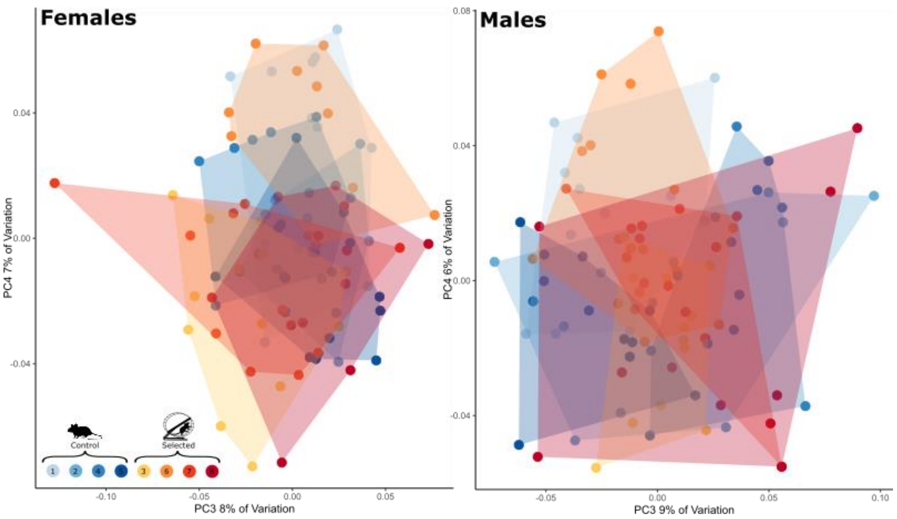


**Supplemental figure 3**. Shape variation associated with PC3 and 4 for males and females.

**Table S4:** ANCOVA results for shape variation associated with individual rib positions for males (Shape~Linetype/Line+Length)

| | Factor | Male | | | | | | |
|---|---|---|---|---|---|---|---|---|
| | | **Df** | **SS** | **MS** | **Rsq** | **F** | **Z** | **Pr(>F)** |
| **Rib 1** | **Linetype** | 1 | 0.010194 | 0.010194 | 0.015422 | 0.595435 | -0.4856 | 0.687831 |
| | **Length** | 24 | 0.121604 | 0.005067 | 0.183976 | 1.077859 | 0.681058 | 0.247575 |
| | **Linetype/line** | 6 | 0.102719 | 0.01712 | 0.155404 | 3.64187 | 5.772445 | 1E-04 |
| | **Residuals** | 68 | 0.319656 | 0.004701 | 0.483611 | | | |
| | **Total** | 99 | 0.660977 | | | | | |
| **Rib 2** | **Linetype** | 1 | 0.001477 | 0.001477 | 0.010029 | 0.228477 | -1.58976 | 0.943506 |
| | **Length** | 24 | 0.022186 | 0.000924 | 0.150619 | 0.844259 | -0.9724 | 0.836716 |
| | **Linetype/line** | 6 | 0.038793 | 0.006466 | 0.263368 | 5.905008 | 5.741633 | 1E-04 |
| | **Residuals** | 68 | 0.074455 | 0.001095 | 0.505476 | | | |
| | **Total** | 99 | 0.147296 | | | | | |
| **Rib 3** | **Linetype** | 1 | 0.003702 | 0.003702 | 0.032368 | 1.235876 | 0.975943 | 0.171883 |
| | **Length** | 24 | 0.021637 | 0.000902 | 0.189169 | 1.130225 | 0.794695 | 0.215778 |
| | **Linetype/line** | 6 | 0.017974 | 0.002996 | 0.157141 | 3.755469 | 4.524808 | 1E-04 |
| | **Residuals** | 68 | 0.054243 | 0.000798 | 0.474224 | | | |
| | **Total** | 99 | 0.114382 | | | | | |
| **Rib 4** | **Linetype** | 1 | 0.010745 | 0.010745 | 0.07206 | 1.865599 | 1.58693 | 0.058794 |
| | **Length** | 24 | 0.020586 | 0.000858 | 0.138064 | 0.743536 | -1.66395 | 0.953005 |
| | **Linetype/line** | 6 | 0.034556 | 0.005759 | 0.231755 | 4.992403 | 5.572606 | 1E-04 |
| | **Residuals** | 68 | 0.078447 | 0.001154 | 0.526111 | | | |
| | **Total** | 99 | 0.149107 | | | | | |
| **Rib 5** | **Linetype** | 1 | 0.043283 | 0.043283 | 0.118148 | 3.721772 | 2.312763 | 0.007699 |
| | **Length** | 24 | 0.051245 | 0.002135 | 0.139883 | 0.757608 | -1.39339 | 0.919008 |
| | **Linetype/line** | 6 | 0.069778 | 0.01163 | 0.190471 | 4.126349 | 4.472556 | 1E-04 |
| | **Residuals** | 68 | 0.19165 | 0.002818 | 0.523142 | | | |
| | **Total** | 99 | 0.366343 | | | | | |

**Table S5:** ANCOVA results for shape variation associated with individual rib positions for females (Shape~Linetype/Line+Length)

| | | Female | | | | | | |
|---|---|---|---|---|---|---|---|---|
| | | **Df** | **SS** | **MS** | **Rsq** | **F** | **Z** | **Pr(>F)** |
| **Rib 1** | **Linetype** | 1 | 0.009014 | 0.009014 | 0.016065 | 0.559768 | -0.63357 | 0.735426 |
| | **Length** | 24 | 0.098583 | 0.004108 | 0.175699 | 0.940929 | -0.62404 | 0.731527 |
| | **Linetype/line** | 6 | 0.09662 | 0.016103 | 0.172199 | 3.68875 | 6.429567 | 1E-04 |
| | **Residuals** | 63 | 0.275028 | 0.004366 | 0.490164 | | | |
| | **Total** | 94 | 0.561093 | | | | | |
| **Rib 2** | **Linetype** | 1 | 0.002867 | 0.002867 | 0.019314 | 0.564188 | -0.16869 | 0.566043 |
| | **Length** | 24 | 0.021202 | 0.000883 | 0.142838 | 0.950045 | -0.26761 | 0.60214 |
| | **Linetype/line** | 6 | 0.030488 | 0.005081 | 0.2054 | 5.464655 | 6.963933 | 1E-04 |
| | **Residuals** | 63 | 0.058581 | 0.00093 | 0.394664 | | | |
| | **Total** | 94 | 0.148432 | | | | | |
| **Rib 3** | **Linetype** | 1 | 0.007781 | 0.007781 | 0.061958 | 2.206195 | 1.780779 | 0.037996 |
| | **Length** | 24 | 0.017189 | 0.000716 | 0.136866 | 0.856557 | -0.97716 | 0.835316 |
| | **Linetype/line** | 6 | 0.021162 | 0.003527 | 0.168501 | 4.218156 | 4.953697 | 1E-04 |
| | **Residuals** | 63 | 0.052679 | 0.000836 | 0.419439 | | | |
| | **Total** | 94 | 0.125593 | | | | | |
| **Rib 4** | **Linetype** | 1 | 0.006305 | 0.006305 | 0.034787 | 1.005544 | 0.703241 | 0.251975 |
| | **Length** | 24 | 0.03168 | 0.00132 | 0.174786 | 1.006746 | 0.085803 | 0.462054 |
| | **Linetype/line** | 6 | 0.037622 | 0.00627 | 0.20757 | 4.782314 | 5.14777 | 1E-04 |
| | **Residuals** | 63 | 0.082602 | 0.001311 | 0.455739 | | | |
| | **Total** | 94 | 0.181249 | | | | | |
| **Rib 5** | **Linetype** | 1 | 0.011616 | 0.011616 | 0.038908 | 1.511521 | 1.20083 | 0.121988 |
| | **Length** | 24 | 0.063494 | 0.002646 | 0.212676 | 1.00751 | 0.087781 | 0.463054 |
| | **Linetype/line** | 6 | 0.04611 | 0.007685 | 0.154446 | 2.926618 | 3.914963 | 1E-04 |
| | **R:esiduals** | 63 | 0.16543 | 0.002626 | 0.554114 | | | |
| | **T;:otal** | 94 | 0.298549 | | | | | |

**Table S6:** ANCOVA results for shape variation associated with individual rib positions for separated by linetype for control males (CTRLshape~Line+Length)

| | | Df | SS | MS | Rsq | F | Z | Pr(>F) |
|---|---|---|---|---|---|---|---|---|
| **Rib1** | **line** | 3 | 0.032 | 0.011 | 0.119 | 2.064 | 2.512 | 0.005 |
| | **length** | 19 | 0.082 | 0.004 | 0.303 | 0.831 | -1.219 | 0.889 |
| | **Residuals** | 25 | 0.130 | 0.005 | 0.480 | | | |
| | **Total** | 47 | 0.271 | | | | | |
| **Rib2** | **line** | 3 | 0.014 | 0.005 | 0.253 | 5.457 | 4.428 | 0.000 |
| | **length** | 19 | 0.014 | 0.001 | 0.248 | 0.843 | -0.860 | 0.805 |
| | **Residuals** | 25 | 0.022 | 0.001 | 0.387 | | | |
| | **Total** | 47 | 0.056 | | | | | |
| **Rib3** | **line** | 3 | 0.008 | 0.003 | 0.160 | 3.467 | 3.249 | 0.000 |
| | **length** | 19 | 0.014 | 0.001 | 0.271 | 0.925 | -0.366 | 0.638 |
| | **Residuals** | 25 | 0.019 | 0.001 | 0.385 | | | |
| | **Total** | 47 | 0.050 | | | | | |
| **Rib4** | **line** | 3 | 0.008 | 0.003 | 0.126 | 2.104 | 1.577 | 0.060 |
| | **length** | 19 | 0.016 | 0.001 | 0.257 | 0.677 | -1.693 | 0.953 |
| | **Residuals** | 25 | 0.030 | 0.001 | 0.500 | | | |
| | **Total** | 47 | 0.060 | | | | | |
| **Rib5** | **line** | 3 | 0.009 | 0.003 | 0.076 | 1.093 | 0.402 | 0.352 |
| | **length** | 19 | 0.036 | 0.002 | 0.314 | 0.709 | -1.235 | 0.889 |
| | **Residuals** | 25 | 0.066 | 0.003 | 0.583 | | | |
| | **Total** | 47 | 0.113 | | | | | |

**Table S7:** ANCOVA results for shape variation associated with individual rib positions for separated by linetype for selected males (Selected shape~Line+Length)

| | | Df | SS | MS | Rsq | F | Z | Pr(>F) |
|---|---|---|---|---|---|---|---|---|

| | | | | | | | | |
|---|---|---|---|---|---|---|---|---|
| **Rib1** | **line** | 3 | 0.032 | 0.011 | 0.119 | 2.064 | 2.512 | 0.005 |
| | **length** | 19 | 0.082 | 0.004 | 0.303 | 0.831 | -1.219 | 0.889 |
| | **Residuals** | 25 | 0.130 | 0.005 | 0.480 | | | |
| | **Total** | 47 | 0.271 | | | | | |
| **Rib2** | **line** | 3 | 0.016 | 0.005 | 0.182 | 4.157 | 3.358 | 0.000 |
| | **length** | 20 | 0.025 | 0.001 | 0.290 | 0.994 | -0.011 | 0.505 |
| | **Residuals** | 28 | 0.036 | 0.001 | 0.408 | | | |
| | **Total** | 51 | 0.088 | | | | | |
| **Rib3** | **line** | 3 | 0.005 | 0.002 | 0.088 | 2.212 | 2.280 | 0.011 |
| | **length** | 20 | 0.023 | 0.001 | 0.405 | 1.529 | 2.047 | 0.018 |
| | **Residuals** | 28 | 0.021 | 0.001 | 0.371 | | | |
| | **Total** | 51 | 0.056 | | | | | |
| **Rib4** | **line** | 3 | 0.016 | 0.005 | 0.181 | 4.233 | 3.646 | 0.0001 |
| | **length** | 20 | 0.019 | 0.001 | 0.221 | 0.776 | -1.130 | 0.871 |
| | **Residuals** | 28 | 0.034 | 0.00122671 | 0.39819316 | | | |
| | **Total** | 51 | 0.086 | | | | | |
| **Rib5** | **line** | 3 | 0.050 | 0.017 | 0.230 | 5.405 | 3.547 | 0.0002 |
| | **length** | 20 | 0.054 | 0.003 | 0.247 | 0.870 | -0.516 | 0.695 |
| | **Residuals** | 28 | 0.087 | 0.003 | 0.397 | | | |
| | **Total** | 51 | 0.220 | | | | | |

**Table S8:** ANCOVA results for shape variation associated with individual rib positions for separated by linetype for control females (Control shape~Line+Length)

| | | **Df** | **SS** | **MS** | **Rsq** | **F** | **Z** | **Pr(>F)** |
|---|---|---|---|---|---|---|---|---|
| **Rib1** | **line** | 3 | 0.041 | 0.014 | 0.169 | 3.147 | 4.076 | 0.0001 |
| | **length** | 19 | 0.078 | 0.004 | 0.326 | 0.955 | -0.366 | 0.647 |
| | **Residuals** | 25 | 0.107 | 0.004 | 0.449 | | | |
| | **Total** | 47 | 0.239 | | | | | |
| **Rib2** | **line** | 3 | 0.011 | 0.004 | 0.221 | 5.920 | 4.950 | 0.0001 |
| | **length** | 19 | 0.023 | 0.001 | 0.434 | 1.831 | 2.219 | 0.007 |
| | **Residuals** | 25 | 0.016 | 0.001 | 0.312 | | | |
| | **Total** | 47 | 0.052 | | | | | |
| **Rib3** | **line** | 3 | 0.009 | 0.003 | 0.172 | 3.931 | 3.161 | 0.0002 |
| | **length** | 19 | 0.017 | 0.001 | 0.325 | 1.171 | 0.770 | 0.224 |
| | **Residuals** | 25 | 0.019 | 0.001 | 0.365 | | | |
| | **Total** | 47 | 0.052 | | | | | |
| **Rib4** | **line** | 3 | 0.017 | 0.006 | 0.199 | 4.191 | 2.848 | 0.0009 |
| | **length** | 19 | 0.021 | 0.001 | 0.243 | 0.806 | -0.780 | 0.781 |
| | **Residuals** | 25 | 0.034 | 0.001 | 0.397 | | | |
| | **Total** | 47 | 0.085 | | | | | |

| | | | | | | | | |
|---|---|---|---|---|---|---|---|---|
| **Rib5** | **line** | 3 | 0.014 | 0.005 | 0.140 | 2.561 | 2.365 | 0.008 |
| | **length** | 19 | 0.040 | 0.002 | 0.414 | 1.192 | 0.744 | 0.230 |
| | **Residuals** | 25 | 0.045 | 0.002 | 0.456 | | | |
| | **Total** | 47 | 0.098 | | | | | |

**Table S9:** ANCOVA results for shape variation associated with individual rib positions for separated by linetype for Selected females (Selected shape~Line+Length)

| | | **Df** | **SS** | **MS** | **Rsq** | **F** | **Z** | **Pr(>F)** |
|---|---|---|---|---|---|---|---|---|
| **Rib1** | **line** | 3 | 0.041 | 0.014 | 0.169 | 3.147 | 4.076 | 1.00E-04 |
| | **length** | 19 | 0.078 | 0.004 | 0.326 | 0.955 | -0.366 | 6.47E-01 |
| | **Residuals** | 25 | 0.107 | 0.004 | 0.449 | | | |
| | **Total** | 47 | 0.239 | | | | | |
| **Rib2** | **line** | 3 | 0.017 | 0.006 | 0.202 | 5.811 | 5.207 | 1.0E-04 |
| | **length** | 19 | 0.018 | 0.001 | 0.217 | 0.985 | -0.073 | 5.2E-01 |
| | **Residuals** | 24 | 0.023 | 0.001 | 0.278 | | | |
| | **Total** | 46 | 0.083 | | | | | |
| **Rib3** | **line** | 3 | 0.008 | 0.003 | 0.142 | 2.701 | 2.429 | 0.005 |
| | **length** | 19 | 0.010 | 0.001 | 0.187 | 0.561 | -2.956 | 0.999 |
| | **Residuals** | 24 | 0.023 | 0.001 | 0.421 | | | |
| | **Total** | 46 | 0.055 | | | | | |
| **Rib4** | **line** | 3 | 0.017 | 0.006 | 0.190 | 3.896 | 3.278 | 0.000 |
| | **length** | 19 | 0.025 | 0.001 | 0.269 | 0.871 | -0.511 | 0.687 |
| | **Residuals** | 24 | 0.036 | 0.00148537 | 0.39031061 | | | |
| | **Total** | 46 | 0.091 | | | | | |
| **Rib5** | **line** | 3 | 0.030 | 0.010 | 0.157 | 2.892 | 2.724 | 0.002 |
| | **length** | 19 | 0.060 | 0.003 | 0.314 | 0.915 | -0.304 | 0.619 |
| | **Residuals** | 24 | 0.083 | 0.003 | 0.433 | | | |
| | **Total** | 46 | 0.193 | | | | | |

**Table S10:** P values of within rib disparity analysis in males

| | | **Control** | | | | | **Selected** | | | | |
|---|---|---|---|---|---|---|---|---|---|---|---|
| | | **Rib 1** | **Rib 2** | **Rib 3** | **Rib 4** | **Rib 5** | **Rib 1** | **Rib 2** | **Rib 3** | **Rib 4** | **Rib 5** |
| **Control** | **Rib 1** | 1 | 0.001 | 0.001 | 0.001 | 0.001 | 0.401 | 0.001 | 0.001 | 0.001 | 0.001 |
| | **Rib 2** | 0.001 | 1 | 0.815 | 0.729 | 0.13 | 0.001 | 0.63 | 0.963 | 0.6 | 0.006 |
| | **Rib 3** | 0.001 | 0.815 | 1 | 0.544 | 0.085 | 0.001 | 0.507 | 0.792 | 0.434 | 0.003 |
| | **Rib 4** | 0.001 | 0.729 | 0.544 | 1 | 0.253 | 0.001 | 0.939 | 0.764 | 0.878 | 0.009 |

| | | | | | | | | | | | |
|---|---|---|---|---|---|---|---|---|---|---|---|
| | **Rib 5** | 0.001 | 0.13 | 0.085 | 0.253 | 1 | 0.001 | 0.291 | 0.147 | 0.342 | 0.125 |
| **Selected** | **Rib 1** | 0.401 | 0.001 | 0.001 | 0.001 | 0.001 | 1 | 0.001 | 0.001 | 0.001 | 0.001 |
| | **Rib 2** | 0.001 | 0.63 | 0.507 | 0.939 | 0.291 | 0.001 | 1 | 0.692 | 0.957 | 0.008 |
| | **Rib 3** | 0.001 | 0.963 | 0.792 | 0.764 | 0.147 | 0.001 | 0.692 | 1 | 0.655 | 0.001 |
| | **Rib 4** | 0.001 | 0.6 | 0.434 | 0.878 | 0.342 | 0.001 | 0.957 | 0.655 | 1 | 0.013 |
| | **Rib 5** | 0.001 | 0.006 | 0.003 | 0.009 | 0.125 | 0.001 | 0.008 | 0.001 | 0.013 | 1 |

**Table S11:** P values of within rib disparity analysis in females

| | | **Control** | | | | | **Selected** | | | | |
|---|---|---|---|---|---|---|---|---|---|---|---|
| | | **Rib 1** | **Rib 2** | **Rib 3** | **Rib 4** | **Rib 5** | **Rib 1** | **Rib 2** | **Rib 3** | **Rib 4** | **Rib 5** |
| **Control** | **Rib 1** | 1 | 0.001 | 0.001 | 0.001 | 0.001 | 0.693 | 0.001 | 0.001 | 0.001 | 0.006 |
| | **Rib 2** | 0.001 | 1 | 0.904 | 0.329 | 0.027 | 0.001 | 0.704 | 0.96 | 0.321 | 0.001 |
| | **Rib 3** | 0.001 | 0.904 | 1 | 0.282 | 0.013 | 0.001 | 0.628 | 0.943 | 0.25 | 0.001 |
| | **Rib 4** | 0.001 | 0.329 | 0.282 | 1 | 0.233 | 0.001 | 0.546 | 0.296 | 0.99 | 0.001 |
| | **Rib 5** | 0.001 | 0.027 | 0.013 | 0.233 | 1 | 0.001 | 0.079 | 0.028 | 0.177 | 0.007 |
| **Selected** | **Rib 1** | 0.693 | 0.001 | 0.001 | 0.001 | 0.001 | 1 | 0.001 | 0.001 | 0.001 | 0.001 |
| | **Rib 2** | 0.001 | 0.704 | 0.628 | 0.546 | 0.079 | 0.001 | 1 | 0.676 | 0.569 | 0.001 |
| | **Rib 3** | 0.001 | 0.96 | 0.943 | 0.296 | 0.028 | 0.001 | 0.676 | 1 | 0.283 | 0.001 |
| | **Rib 4** | 0.001 | 0.321 | 0.25 | 0.99 | 0.177 | 0.001 | 0.569 | 0.283 | 1 | 0.001 |
| | **Rib 5** | 0.006 | 0.001 | 0.001 | 0.001 | 0.007 | 0.001 | 0.001 | 0.001 | 0.001 | 1 |

**Table S12:** P values of within rib integration analysis in males

| **Selected** | | | | **Control** | | | | | | |
|---|---|---|---|---|---|---|---|---|---|---|
| **Rib 2** | **Rib 3** | **Rib 4** | **Rib 5** | **Rib 1** | **Rib 2** | **Rib 3** | **Rib 4** | **Rib 5** | | |
| 0.128 | 0.101 | 0.077 | 0.059 | 0.057 | 0.340 | 0.212 | 0.040 | 0.019 | **Selected** | **Rib 2** |
| | 0.443 | 0.386 | 0.335 | 0.004 | 0.064 | 0.378 | 0.261 | 0.168 | | **Rib 3** |
| | | 0.441 | 0.388 | 0.002 | 0.048 | 0.326 | 0.308 | 0.205 | | **Rib 4** |
| | | | 0.446 | 0.001 | 0.035 | 0.276 | 0.361 | 0.249 | | **Rib 5** |
| | | | | 0.001 | 0.026 | 0.233 | 0.412 | 0.293 | | **Rib 1** |

| | | | | | | | | | | |
|---|---|---|---|---|---|---|---|---|---|---|
| | | | | | 0.127 | 0.010 | 0.001 | 0.000 | Control | **Rib 2** |
| | | | | | | 0.117 | 0.017 | 0.007 | | **Rib 3** |
| | | | | | | | 0.176 | 0.106 | | **Rib 4** |
| | | | | | | | | 0.376 | | **Rib 5** |

**Table S13:** P values of within rib integration analysis in females

| **Selected** | | | | **Control** | | | | | | |
|---|---|---|---|---|---|---|---|---|---|---|
| **Rib 2** | **Rib 3** | **Rib 4** | **Rib 5** | **Rib 1** | **Rib 2** | **Rib 3** | **Rib 4** | **Rib 5** | | |
| 0.015 | 0.004 | 0.006 | 0.017 | 0.318 | 0.021 | 0.004 | 0.000 | 0.005 | **Selected** | **Rib 2** |
| | 0.322 | 0.361 | 0.473 | 0.004 | 0.433 | 0.310 | 0.022 | 0.359 | | **Rib 3** |
| | | 0.457 | 0.298 | 0.001 | 0.263 | 0.488 | 0.060 | 0.458 | | **Rib 4** |
| | | | 0.336 | 0.001 | 0.299 | 0.446 | 0.048 | 0.499 | | **Rib 5** |
| | | | | 0.005 | 0.459 | 0.287 | 0.018 | 0.334 | | **Rib 1** |
| | | | | | 0.006 | 0.001 | 0.000 | 0.001 | **Control** | **Rib 2** |
| | | | | | | 0.252 | 0.014 | 0.297 | | **Rib 3** |
| | | | | | | | 0.062 | 0.446 | | **Rib 4** |
| | | | | | | | | 0.047 | | **Rib 5** |
